# Img2EEG: A Scalable and Interpretable Encoding Framework for Simulating Human EEG Responses to Visual Inputs

**DOI:** 10.64898/2026.09.16.751610

**Authors:** Zitong Lu, Julie Golomb

## Abstract

Understanding how visual information processing unfolds over time requires models that not only predict neural responses but also expose the representations that support them and generalize beyond sampled stimulus spaces. Here we introduce Img2EEG, a participant-specific image-to-EEG encoding framework that integrates hierarchical visual and semantic representations to generate temporally resolved multichannel EEG responses. Trained on THINGS EEG2, Img2EEG generalized to unseen images while preserving stimulus-specific and participant-specific response structure. Controlled perturbations of internal representations and visual inputs revealed distinct temporally structured contributions of visual and semantic information, and in silico experiments reproduced classic human neural responses, such as the face-sensitive N170, while enabling targeted representational interventions. Scaling Img2EEG to 1.28 million ImageNet images produced over 12 million synthetic EEG responses that supported cross-dataset visual reconstruction and improved the behavioral alignment of an artificial vision model. Img2EEG provides an interpretable and scalable framework for experimentally manipulable modeling of visual neural dynamics.

## Introduction

Computational encoding models provide a powerful framework for formalizing how sensory inputs are transformed into neural responses (Kriegeskorte & Douglas, 2019; Naselaris et al., 2011). In vision, encoding models have been widely used with functional magnetic resonance imaging (fMRI) to map rich image representations from artificial neural networks onto spatially resolved cortical activity (Gifford et al., 2023; Huth et al., 2012, 2016; Naselaris et al., 2015). Yet visual processing unfolds over tens to hundreds of milliseconds, and understanding this temporal evolution requires models that can predict neural dynamics at a correspondingly fine timescale. Electroencephalography (EEG) offers the temporal resolution needed for this goal, but building image-to-EEG encoding models remains challenging because visually evoked EEG responses are high-dimensional, noisy, strongly time-dependent, and vary substantially across individuals. Recent work has begun to address this problem, showing that visual representations can be mapped onto temporally resolved EEG responses and used to generate synthetic electrophysiological responses to unseen stimuli (Gifford et al., 2022; Santos-Mayo et al., 2026). The broader challenge is to determine **whether such models can become more than predictors of brain activity – whether they can serve as interpretable computational models for probing how visual information unfolds over time in the human brain.**

Accurate prediction of EEG responses alone provides limited insight into the computations that give rise to those responses (Kriegeskorte & Douglas, 2019; Yamins & DiCarlo, 2016). First, high similarity between predicted and measured EEG does not necessarily indicate that a model captures the fine-grained neural information distinguishing one visual input from another, because stereotyped response dynamics shared across stimuli can contribute substantially to overall prediction accuracy. Second, prediction-oriented image-to-EEG models do not necessarily expose which forms of stimulus information support different portions of the predicted neural response. Existing approaches have largely emphasized accurate response generation from compact visual or conceptual representations, rather than enabling controlled intervention on distinct representational components. For example, Concept2Brain maps a CLIP-derived conceptual representation onto a learned electrophysiological latent space and is primarily designed to reproduce characteristic EEG waveforms and topographies rather than to explain the computations underlying them (Santos-Mayo et al., 2026). An interpretable image-to-EEG model should therefore not only preserve stimulus-specific neural information, but also support controlled manipulation of distinct representations and generate testable in silico predictions about established neural phenomena. Ideally, such a model should be able to recapitulate canonical EEG effects and use targeted perturbations to identify the representational factors required for their emergence within the model.

Yet even a highly interpretable model remains constrained by the limited scale of empirical human EEG data. The ability to scale image-to-EEG modeling therefore depends not only on generating responses to new stimuli, but also on learning neural mappings from a sufficiently broad and diverse visual space. For example, Concept2Brain was trained on only 360 unique naturalistic images, with the authors noting the limited size of the model’s current concept space (Santos-Mayo et al., 2026). Large-scale datasets such as THINGS EEG2 (Gifford et al., 2022), which build on the broad THINGS object-concept space (Hebart et al., 2019; Stoinski et al., 2023), provide an important opportunity in this regard by sampling thousands of natural images spanning a broad range of object concepts, allowing image-to-EEG mappings to be learned from a substantially richer visual stimulus space and evaluated on entirely unseen images and concepts. A framework that can reliably capture such mappings could then extend model-based neural predictions to much larger image spaces, opening the possibility of studying human visual dynamics at a scale that would be difficult to achieve through empirical recording alone.

Here, we introduce Img2EEG, a scalable and interpretable encoding framework for simulating human EEG from visual inputs. Img2EEG learns participant-specific mappings from natural images to temporally resolved multichannel EEG responses by integrating hierarchical visual and higher-level semantic representations. Using the THINGS EEG2 dataset, we first establish that these mappings generalize to unseen images and concepts while preserving both stimulus-specific and participant-specific neural information. We then use targeted perturbations to transform the model from a predictor into an experimentally manipulable in silico system, revealing when different types of visual and semantic information are required to account for human EEG dynamics. We next test whether this framework can reproduce established visual EEG phenomena. We then use targeted perturbations to probe which internal representations contribute to the model’s reproduction of these effects. Finally, we scale the learned mappings to more than 1.2 million ImageNet images and ask whether the resulting synthetic EEG remains informative when transferred back to independently measured human EEG and brain-aligned artificial vision models. Together, these analyses establish Img2EEG as a framework for moving from finite empirical measurements toward predictive, interpretable and scalable models of human visual dynamics.

## Results

### Img2EEG predicts individualized and image-specific EEG responses to unseen visual inputs

We first asked whether a participant-specific image-to-EEG framework could faithfully capture stimulus-specific human visual EEG responses and support subsequent in silico experimentation. We therefore developed Img2EEG, which combines complementary visual and semantic representations to predict the spatiotemporal EEG response evoked by an image (Figure 1A). The hierarchical visual encoder incorporates representations from multiple stages (layer V1, V2, and V4) of CORnet-S (Kubilius et al., 2018, 2019), capturing multiple levels of visual information, whereas the semantic encoder combines a CLIP visual embedding (Radford et al., 2021), a GloVe-based object-concept representation (Pennington et al., 2014), and an image-description representation obtained by embedding BLIP2-generated captions (J. Li et al., 2023) with MPNet (K. Song et al., 2020). The outputs of the two encoding pathways are projected and integrated through a series of learned layers, with the final layer generating an 850-dimensional response corresponding to 17 EEG channels across 50 time points.

**Figure 1.**
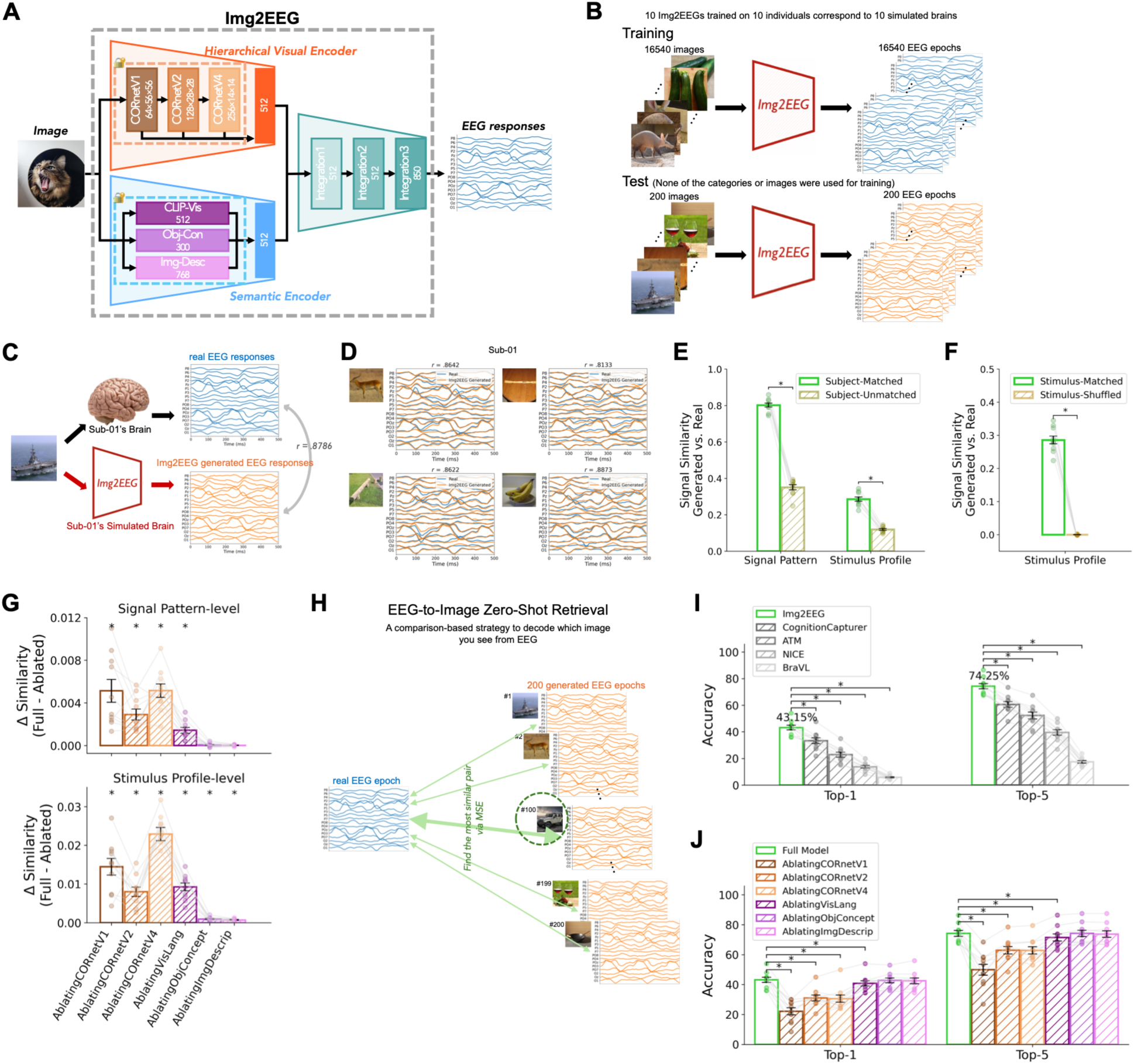
Img2EEG generates individualized and image-specific EEG responses to unseen visual inputs. (A) Architecture of Img2EEG. Each input image is represented by two complementary encoding pathways. The hierarchical visual encoder extracts representations from multiple stages (including layer V1, V2, and V4) of CORnet-S corresponding to multiple levels of visual processing. The semantic encoder incorporates three higher-level representations: a CLIP visual embedding (CLIP-Vis), an object-concept representation (Obj-Con) derived from GloVe, and an image-description representation (Img-Desc) obtained by first generating an image caption with BLIP-2 and then embedding the caption with MPNet. Representations within each pathway are projected into a common feature space and subsequently combined through a series of integration layers. The final integration layer outputs 850 values corresponding to the complete predicted EEG response (17 channels × 50 time points), thereby mapping each visual input directly onto a multichannel, temporally resolved EEG epoch. (B) Training and test scheme. Ten participant-specific Img2EEG models are trained separately using 16,540 images and their corresponding EEG epochs from the THINGS EEG2 training set. Model performance was evaluated on 200 held-out images and corresponding EEG epochs; neither the test images nor their object concepts / categories were included during training. Each participant-specific model therefore serves as an individualized simulated brain. (C) Example comparison between the real EEG responses of Sub-01 and the EEG responses generated by the corresponding Img2EEG model for an unseen test image. (D) Additional examples illustrating the correspondence between real and generated EEG responses across unseen images for Sub-01. (E) Similarity between real and generated EEG responses when the Img2EEG model and the participant were matched or unmatched, quantified at both the signal-pattern and stimulus-profile levels. (F) Stimulus specificity of stimulus-profile similarity. Stimulus-profile similarity was compared between the original stimulus-matched analysis and a stimulus-shuffled control in which image correspondence between generated and empirical EEG responses was randomly permuted across the 200 held-out test images. Higher similarity in the matched condition indicates preservation of image-specific neural response structure. (G) Reduction in prediction similarity following ablation of individual visual or semantic components in Img2EEG, evaluated at both the signal-pattern and stimulus-profile levels. Positive values indicate lower similarity in the ablated than in the full model. (H) Schematic of the EEG-to-image zero-shot retrieval procedure using Img2EEG. For each real EEG epoch, Img2EEG generates candidate EEG epochs for all 200 held-out images, and the image whose generated epoch has the smallest mean-squared error (MSE) relative to the real epoch is selected as the predicted stimulus. (I) Top-1 and Top-5 zero-shot image-retrieval performance of Img2EEG compared with existing EEG-to-image decoding models (J) Effects of ablating individual Img2EEG components on Top-1 and Top-5 zero-shot retrieval performance. Points denote individual participants; Error bars indicate SEM; Asterisks indicate statistically significant differences (*p* < .05, FDR-corrected). Detailed procedures are described in the Methods.

We trained a separate Img2EEG model for each of the ten participants in the THINGS EEG2 dataset (Figure 1B). Each model was trained using EEG responses to 16,540 natural images and evaluated on an independent set of 200 images. Importantly, neither the test images nor their object concepts were included during model training, providing a stringent test of generalization to unseen visual content. Thus, rather than learning a single population-average mapping, the framework yielded ten individualized image-to-EEG models that could be treated as participant-specific simulated response systems.

Img2EEG generated EEG responses that closely resembled the corresponding real responses for unseen images (Figure 1C–D). Signal-pattern similarity measured correspondence between the complete generated and empirical spatiotemporal EEG patterns for each image, while stimulus-profile similarity measured whether the model preserved relative response variation across the 200 test images at each channel and time point. Both forms of correspondence were participant-specific: responses generated by a participant’s own Img2EEG model were significantly more similar to that participant’s real EEG than responses generated by models trained on other participants (Figure 1E; subject-matched vs. subject-unmatched: 0.8018 vs. 0.3509 for signal pattern; 0.2858 vs. 0.1191 for stimulus profile). Consistent with the participant-specificity of the predicted EEG responses, representational similarity across individualized models also varied by processing stage, with semantic representations showing greater cross-participant consistency than hierarchical visual and integration representations (Figure S1).

We additionally asked whether the stimulus-profile correspondence specifically depended on correct image identity. We therefore repeated the analysis after randomly shuffling the correspondence between generated and empirical responses across the 200 test images. Stimulus-profile similarity was substantially higher when stimulus identity was preserved than after shuffling (Figure 1F; stimulus-matched vs. stimulus-shuffled: 0.2858 vs. -0.0001), demonstrating that Img2EEG captured systematic image-specific variation rather than only response structure shared across stimuli.

We next asked whether the different representational components of Img2EEG made significant and distinct contributions to EEG prediction. We therefore selectively ablated each visual or semantic component and quantified the resulting reduction in generation performance at both the signal-pattern and stimulus-profile levels (Figure 1G). Ablating representations from the hierarchical visual pathway and higher-level visual information from CLIP reduced prediction performance at both levels, whereas the object-concept and image-description representations selectively affected stimulus-profile similarity. This dissociation suggests that visually grounded representations contribute broadly to reconstructing the spatiotemporal response, whereas explicitly semantic information contributes more selectively to preserving stimulus-dependent variation. Together, these results indicate that successful EEG generation relies on complementary information distributed across the visual and semantic representations rather than on a single dominant representation.

High overall signal similarity, however, does not by itself establish that generated EEG responses contain sufficient information to distinguish among individual images. The stimulus-profile analysis provided evidence that Img2EEG captured systematic image-dependent variation across stimuli, but it did not test whether this information was sufficiently discriminative to identify a specific unseen image among competing alternatives. We therefore tested this more directly using a zero-shot retrieval task on THINGS EEG2 test set using a comparison-based strategy (Figure 1H). For each real EEG response in the held-out test set, we generated candidate EEG responses for all 200 unseen images and identified the candidate whose generated response most closely matched the real EEG. Because none of these images or their object concepts had been used to train Img2EEG, successful retrieval required the model to generate sufficiently distinctive EEG predictions for novel visual inputs to recover stimulus identity.

Img2EEG supported robust zero-shot retrieval. Img2EEG achieved higher Top-1 and Top-5 identification accuracy than the previously reported benchmark values from the comparison EEG-to-image decoding models (Du et al., 2023; D. Li et al., 2024; Y. Song et al., 2024; Zhang et al., 2025) (Figure 1I). This result provides a stronger test of stimulus specificity than overall waveform correspondence: the generated response for the correct image was sufficiently distinct from responses generated for competing images to recover stimulus identity directly in EEG space.

To determine which representations supported this ability, we repeated the retrieval analysis after selectively ablating individual Img2EEG components (Figure 1J). Retrieval performance was most strongly disrupted by removing representations from the visual features, whereas ablation of semantic representations produced substantially smaller effects. Thus, although both visual and semantic information contributed to accurate EEG generation, fine-grained identification of unseen images depended predominantly on visual representational information.

Together, these results establish that Img2EEG does more than reproduce the average structure of visually evoked EEG. The framework generalizes to unseen visual inputs while preserving both stimulus-specific information and participant-specific response structure. This combination of predictive fidelity and specificity provides the prerequisite for treating individualized Img2EEG models as in silico experimental systems whose internal representations can subsequently be manipulated to ask what information supports different components of the human EEG response.

### Interpreting Img2EEG reveals temporally resolved contributions of visual features and representations

Having established the predictive fidelity and specificity of Img2EEG, and identified the overall contributions of its visual and semantic components to EEG generation, we next asked when these contributions emerged over time. Because the model exposes separable visual and semantic representations that can be selectively manipulated while leaving the remaining system unchanged, it provides an opportunity to resolve when different types of information are required for the model to account for human EEG dynamics.

We first performed time-resolved internal representation ablations across the hierarchical visual and semantic encoders (Figure 2A). For each representational component, we removed image-specific information by replacing the representation evoked by each test image with the mean embedding computed across the training set, while keeping all other representations and model parameters unchanged. We then propagated the perturbed representation through the remaining Img2EEG architecture and compared the resulting EEG response with the corresponding real response at each time point (Figure 2B). The loss of prediction similarity relative to the intact model provided a time-resolved measure of the extent to which image-specific information carried by that representation was required to explain the measured EEG response.

**Figure 2.**
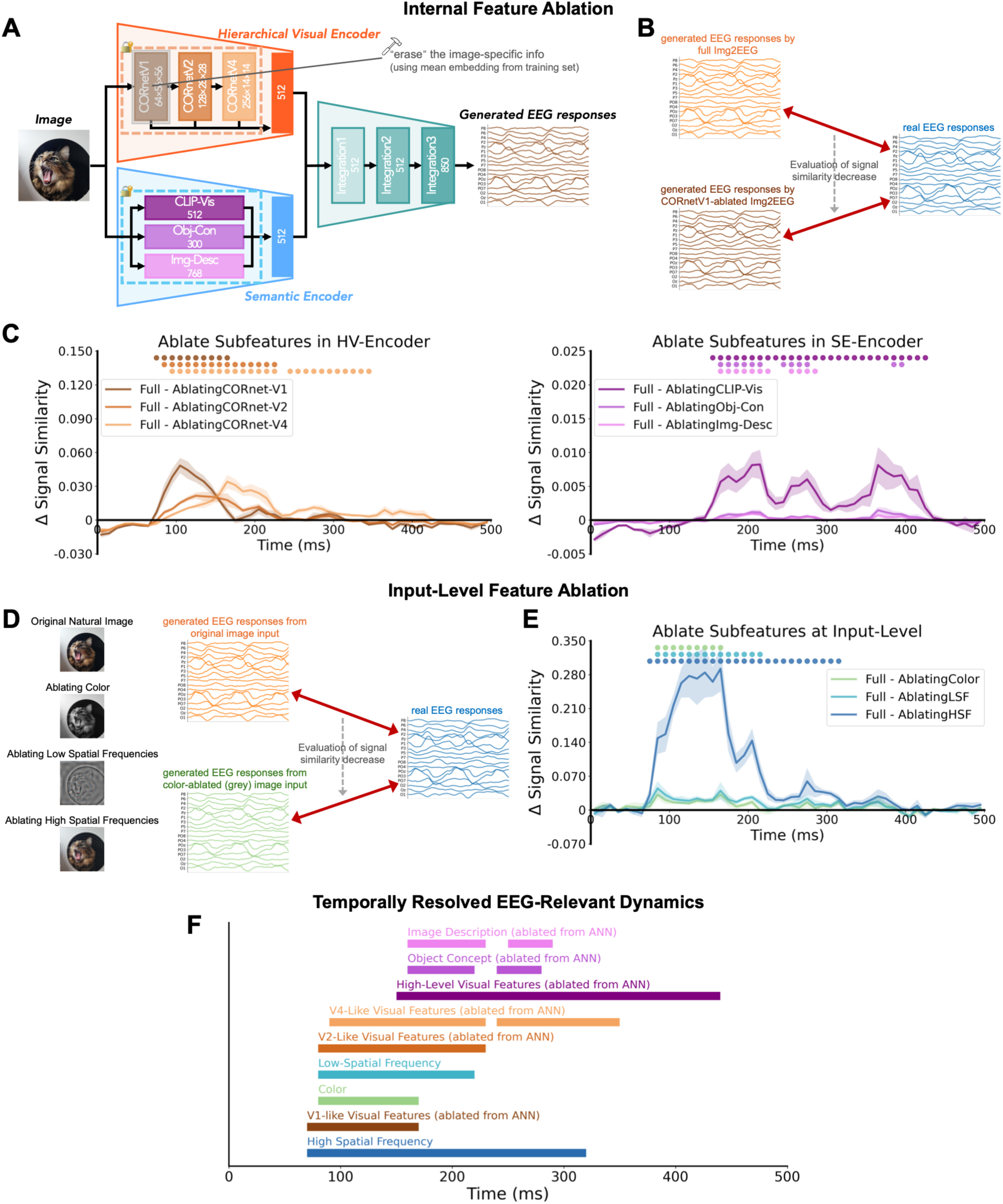
Controlled perturbations of Img2EEG reveal temporally resolved contributions of different features and representations to human EEG dynamics. (A) Schematic of internal feature ablation. To assess the contribution of individual feature components within Img2EEG, image-specific information in a selected feature representation was removed by replacing its embedding with the mean embedding computed across the training set, while all other model inputs and parameters were kept unchanged. The ablated representation was then propagated through the trained integration layers to generate a perturbed EEG response. (B) Evaluation of the effect of internal feature ablation. EEG responses generated by the full and feature-ablated Img2EEG models were separately compared with the corresponding real EEG responses at each time point; the reduction in prediction similarity after ablation was used to quantify the contribution of the removed representation. (C) Time-resolved effects of internal feature ablation. Changes in EEG prediction similarity are shown following removal of representations from the hierarchical visual encoder (left; CORnet-V1, CORnet-V2, and CORnet-V4) and semantic encoder (right; CLIP-Vis, Obj-Con, and Img-Desc). Positive values indicate that removing the targeted image-specific representation reduced correspondence between generated and real EEG responses. (D) Schematic of input-level feature ablation. Visual properties of the input images were selectively manipulated, including removal of color, low spatial frequencies, or high spatial frequencies, and the resulting images were passed through the otherwise unchanged Img2EEG model. The EEG responses generated from the original and feature-ablated images were separately compared with the real EEG responses. (E) Time-resolved reduction in EEG prediction similarity following input-level removal of color, low spatial frequencies (LSF) or high spatial frequencies (HSF). (F) Summary of temporally resolved EEG-relevant information identified by the internal- and input-level perturbation analyses. Horizontal bars indicate time intervals during which removal of each visual or semantic feature significantly reduced correspondence between generated and real EEG responses, illustrating the temporal profiles with which different forms of stimulus information contribute to Img2EEG predictions. Lines and shaded regions in C and E denote the mean and SEM across participants, respectively. Colored markers above the time courses denote statistically significant time points (*p* < .05, cluster-based corrected). Detailed procedures are described in the Methods.

The resulting perturbation profiles revealed temporally differentiated contributions across representational levels (Figure 2C). Ablation of early hierarchical visual representations preferentially reduced prediction accuracy during relatively early portions of the visually evoked response, whereas representations from progressively higher stages contributed over distinct and partly overlapping temporal intervals. Higher-level semantic representations showed later and more selective effects. Thus, the representations supporting Img2EEG predictions were not uniformly informative throughout the response; instead, different forms of visual and semantic information made temporally structured contributions to explaining human EEG dynamics. Importantly, these effects reflect the time periods during which information contained in a model representation was necessary for maintaining accurate EEG predictions, rather than the activation time of the corresponding artificial network layer itself.

We next asked whether temporally specific contributions could also be recovered by manipulating interpretable properties of the visual stimulus itself. Rather than intervening on internal model representations, we selectively removed color, low spatial frequencies or high spatial frequencies from each input image and passed the manipulated images through the otherwise unchanged Img2EEG model (Figure 2D). We again quantified the reduction in correspondence with real EEG relative to predictions generated from the original images. These input-level perturbations produced distinct temporal profiles (Figure 2E), demonstrating that the contribution of physically interpretable visual features to EEG prediction could be localized in time. Removing color, low-spatial-frequency, and high-spatial-frequency information each reduced correspondence between generated and real EEG responses over specific post-stimulus periods, with partially overlapping temporal profiles across features.

Together, the internal- and input-level perturbation analyses yielded a temporally resolved map of EEG-relevant information (Figure 2F). Critically, the two approaches interrogate Img2EEG at complementary levels: internal ablations isolate information represented at different stages of the model, whereas stimulus-level perturbations manipulate properties of the visual input itself. Together, these complementary perturbation analyses show that Img2EEG can be used not only to predict an EEG response, but also to experimentally manipulate the information supporting that prediction. Img2EEG therefore provides an in silico model in which hypotheses about the temporal contribution of specific stimulus features and internal representations can be tested through controlled intervention.

### Img2EEG recapitulates canonical visual EEG phenomena and probes their computational basis

If Img2EEG is to serve as an in silico model of human visual electrophysiology, it should capture not only point-by-point EEG responses but also established neural phenomena that emerge across controlled stimulus contrasts. We therefore asked whether the individualized models could reproduce canonical visual EEG effects and whether targeted interventions could reveal the model-internal representations supporting them. As an initial test of this generalization, we found that Img2EEG reproduced an early lateralized response to unilateral visual stimuli (Figure S2), consistent with the spatial organization of early visual EEG responses (Luck et al., 1990; Mangun & Hillyard, 1990). We then focused on more complex visual EEG phenomena. As a case study, we asked whether Img2EEG might naturally reproduce the N170, a canonical face-sensitive ERP component marked by an enhanced negative potential appearing over posterior electrodes approximately 170ms after stimulus onset, when participants view face images compared to non-face objects (Hinojosa et al., 2015; Mercure et al., 2011; Rossion & Jacques, 2012). We conducted an *in silico* experiment by presenting novel images of faces and objects (not part of the training dataset) to each of the ten individualized Img2EEG models (Figure 3A). Strikingly, the resulting simulated ERPs reproduced the face-selective response in the correct temporal range and posterior electrode sites (Figure 3A). As with a true N170 response, this effect was spatially selective: its magnitude varied systematically across the electrode array, with stronger effects at posterior right-lateralized sites, particularly P8 and PO8 (Bentin et al., 1996; Proverbio, 2021; Rossion & Jacques, 2012) (Figure 3B). Thus, Img2EEG automatically recapitulated both the temporal and spatial characteristics of this canonical face-sensitive EEG phenomenon using images that were not part of the EEG training set.

**Figure 3.**
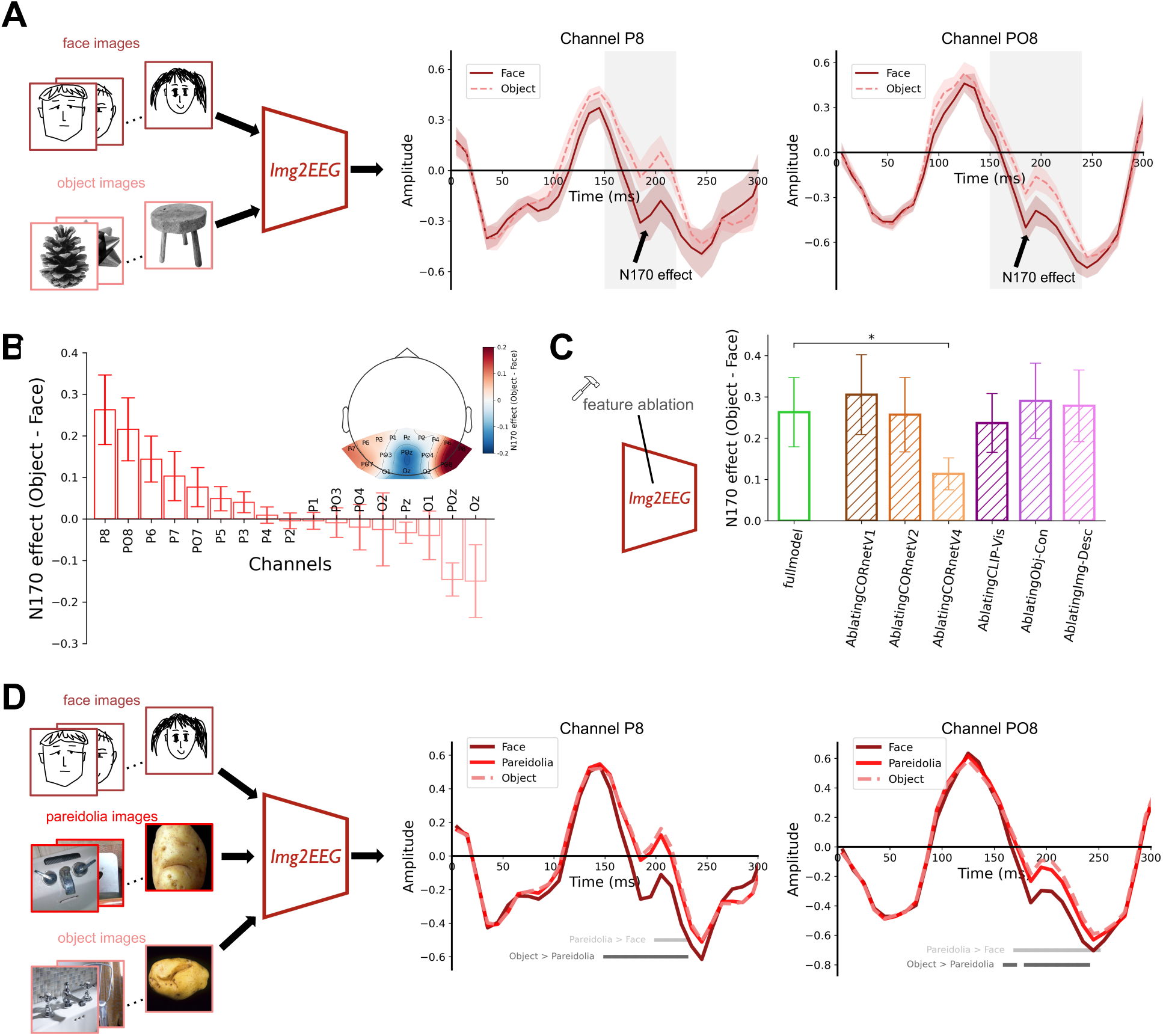
In silico experiments with Img2EEG recapitulate canonical face N170 effects and probe their model-internal representational basis. (A) Schematic of the in silico face-selectivity experiment and simulated face- and object-evoked responses at posterior channels P8 and PO8. Face and non-face object images that were not used for EEG-model training were entered into the individualized Img2EEG models to generate simulated event-related responses. The difference between conditions recapitulated the characteristic face-selective N170 effect in the expected post-stimulus time window. Shaded regions indicate SEM across participants; grey horizontal lines denote time intervals with statistically significant face–object differences (*p* < .05, cluster-based corrected). (B) N170 effect across EEG channels, quantified as the difference in N170 amplitude between object- and face-evoked responses in the 170-200 ms time window. The effect varied systematically across channels rather than emerging uniformly across the scalp. (C) Effect of selectively ablating individual Img2EEG representations on the simulated N170 effect. The face-selective N170 effect is shown for the full model and after ablation of different feature components within Img2EEG. Ablation of the CORnet-V4 representation produced the largest reduction which was significant in the simulated N170 effect, indicating a particularly strong contribution of mid-level visual information to the model’s reproduction of face-selective EEG responses. Bars indicate mean ± SEM across participants; asterisks denote significant differences from the full model (*p* < .05, FDR-corrected) (D) Schematic of the in silico pareidolia experiment and simulated face-, pareidolia-, and object-evoked responses at posterior channels P8 and PO8. Pareidolia stimuli elicited responses intermediate between those evoked by real faces and matched non-face objects, consistent with graded sensitivity to face-like visual structure. SEM shading is omitted from the ERP traces here for visual clarity, allowing the three highly overlapping condition-specific time courses to be distinguished more easily. Detailed procedures are described in the Methods.

Moreover, direct exposure to face-containing EEG training examples was not necessary for the emergence of the simulated N170 effect. After removing all training images containing faces from the THINGS EEG2 training set and trained a new set of participant-specific Img2EEG models using only the remaining images,these non-face-trained Img2EEG models continued to reproduce the characteristic face-versus-object N170 effect and its posterior spatial distribution (Figure S3). This result indicates that direct face-specific EEG supervision is not required for Img2EEG to generate a face-sensitive N170-like response. Instead, the effect shows that a face-sensitive response can be generated from an EEG mapping learned across the broader object space, without direct face-specific EEG supervision, potentially by exploiting visual representational dimensions that distinguish face-like structure from other objects. Because the visual feature extractors themselves were pretrained independently of the EEG data, however, this result does not imply that the underlying artificial representations developed without prior exposure to faces.

Having established that Img2EEG can reproduce the N170, we next exploited a key advantage of the model over conventional EEG experiments: the ability to intervene directly on its internal representations. We selectively ablated individual components of the hierarchical visual and semantic encoders and recomputed the face–object N170 difference (Figure 3C). The results showed that removing the CORnet V4-like representation produced the largest reduction, which was significant in the simulated effect. This pattern suggests that intermediate-level visual information represented within the hierarchical visual pathway is particularly important within Img2EEG for reproducing the face-selective EEG effect.

Having established that Img2EEG recapitulates the canonical face-sensitive N170, we asked whether the model could also reproduce a more subtle and recently characterized phenomenon of face processing: face pareidolia, which occurs when inanimate objects evoke a compelling illusory perception of a face. To provide a more graded test of face-like processing, we presented Img2EEG with the highly controlled face-pareidolia stimulus set (Wardle et al., 2020), comprising real faces, inanimate objects containing illusory faces, and matched non-face objects. In Img2EEG, pareidolia images likewise elicited an N170 response intermediate between those evoked by real faces and ordinary objects at posterior electrodes (Figure 3D) consistent with previous N170 studies. This graded response suggests that the simulated N170 is sensitive not simply to discrete category labels, but to visual structure associated with face-like appearance. More broadly, this type of in silico replication provides a starting point for asking new questions that are difficult to address from scalp EEG alone – for example, whether pareidolia and real faces depend on the same internal representations, or whether different representational components selectively support illusory versus veridical face processing.

Together, these in silico experiments extend the validation of Img2EEG beyond point-by-point prediction of measured EEG. The framework recapitulates established spatial and category-sensitive electrophysiological phenomena, generalizes the N170 effect to graded face-like stimuli, and reproduces it even without direct face-containing EEG training examples. Combined with targeted representation ablations, these findings show that Img2EEG can move beyond reproducing known EEG effects to generate model-based hypotheses about the representational dependencies sufficient for their emergence. These experiments therefore illustrate how an image-to-EEG predictor can be converted into a hypothesis-generating in silico model of visual electrophysiology.

### Scaling Img2EEG creates an ImageNet-scale synthetic EEG resource for visual reconstruction and NeuroAI

Having established Img2EEG as a predictive and experimentally manipulable in silico model, we finally asked whether the learned image-to-EEG mappings could be extended to a visual space far larger than could be sampled experimentally. To do so, we created ImageNet-SimEEG by applying each participant-specific Img2EEG model to the full ImageNet image set of 1.28 million images (Deng et al., 2009). Across all images and ten individualized models, this yielded more than 12 million model-derived EEG epochs (Figure 4A). We then asked whether this large synthetic neural response space could be used for two downstream applications that go beyond EEG generation itself: reconstructing visual content from independently measured human EEG, and providing large-scale neural supervision for brain-aligned artificial vision models.

**Figure 4.**
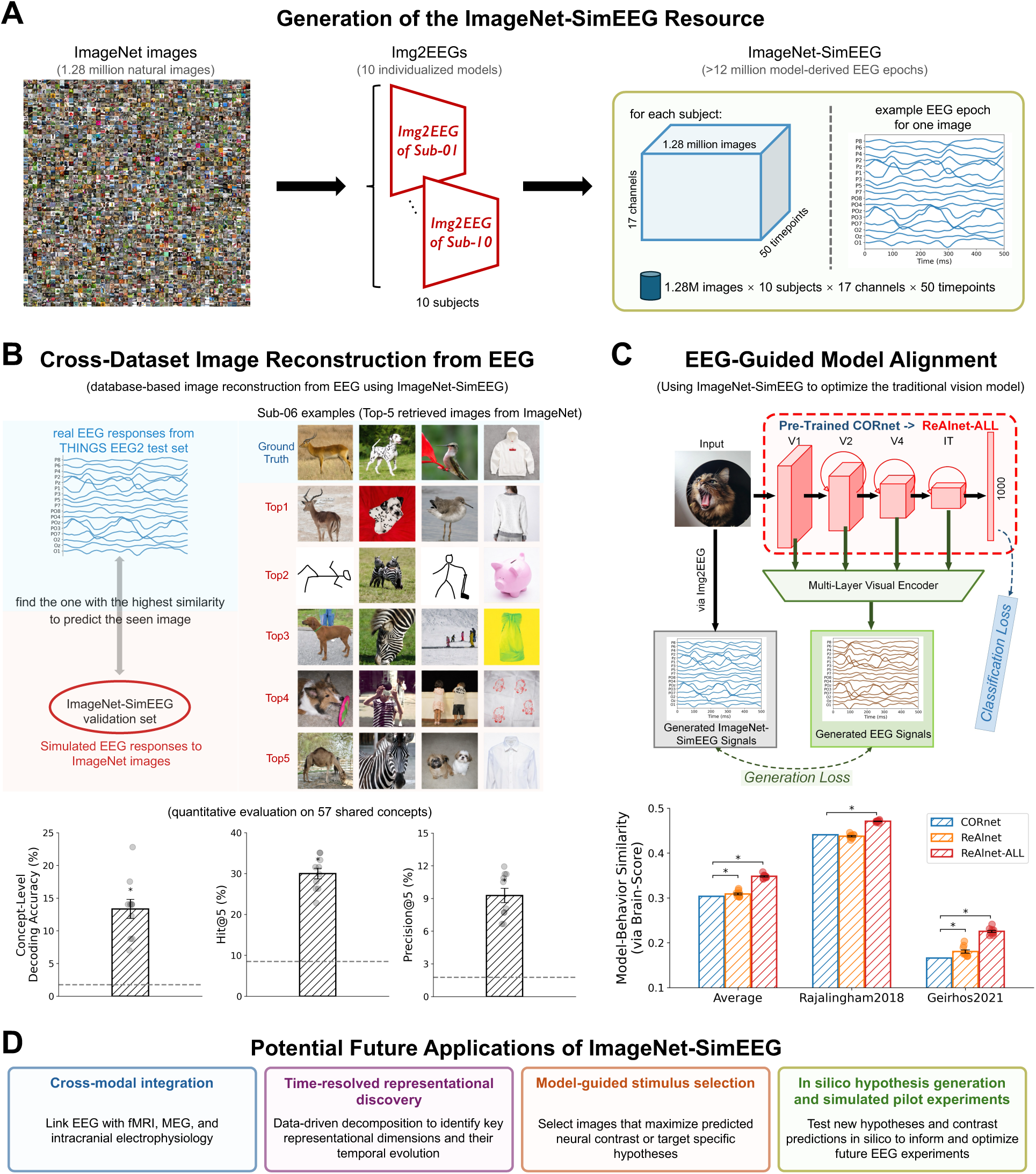
Scaling Img2EEG creates an ImageNet-scale synthetic EEG resource for neural decoding and NeuroAI. (A) Generation of the ImageNet-SimEEG resource. More than 1.28 million ImageNet images were passed through 10 individualized Img2EEG models to generate over 12 million model-derived EEG epochs. For each simulated participant, the resulting resource spans 1.28 million images, 17 EEG channels and 50 time points, yielding a structured image × participant × channel × time response space. (B) Database-based image reconstruction from EEG using ImageNet-SimEEG. Real EEG responses from THINGS EEG2 test set were used as queries against the ImageNet-SimEEG response space. For each real EEG response, ImageNet validation set images were ranked according to the similarity between their simulated EEG responses and the measured EEG pattern, yielding candidate visual reconstructions from a large set of images that had never been viewed during the THINGS EEG2 experiment. Example Top-5 reconstructions are shown for Sub-06. Quantitative performance was evaluated on 57 object concepts shared between THINGS EEG2 and ImageNet using concept-level decoding accuracy, Hit@5 and Precision@5 (Details are described in Methods). (C) EEG-guided model-brain alignment using ImageNet-SimEEG. ImageNet-SimEEG was used as a large-scale neural target to optimize a pretrained CORnet-S vision model, yielding ReAlnet-ALL. During optimization, model-derived EEG responses generated from intermediate visual features were aligned with the corresponding ImageNet-SimEEG responses while retaining the original image-classification objective. The resulting model showed improved model-behavior alignment relative to the pretrained CORnet and real EEG-optimized ReAlnet on Brain-Score platform. (D) Potential future applications of ImageNet-SimEEG. Large-scale synthetic EEG may support cross-modal integration with fMRI, MEG and intracranial electrophysiology; data-driven discovery of time-resolved representational dimensions; model-guided stimulus selection for targeted experiments; and in silico hypothesis generation and simulated pilot experiments to inform future empirical EEG studies. Points in B and C denote individual participants or models. Error bars indicate SEM. Dashed horizontal lines in B indicate chance-level performance. Asterisks denote statistically significant differences (*p* < .05, FDR-corrected). Detailed procedures are described in the Methods.

We found that ImageNet-SimEEG supported cross-dataset reconstruction of visual content from independently measured human EEG. For each empirically recorded human EEG response from the THINGS EEG2 test set, we attempted to reconstruct what the participant was viewing (Figure 4B). We calculated the similarity between each empirical EEG response and the ImageNet-SimEEG pool of simulated responses generated for ImageNet images and ranked the ImageNet candidate images according to their similarity. This procedure therefore used ImageNet-SimEEG as a large model-derived neural lookup space to infer visual content from brain responses without training a separate EEG-to-image decoder. Quantitative evaluation was performed at the concept level for the 57 object concepts shared between THINGS EEG2 and ImageNet. Reconstruction performance exceeded chance for across concept-level decoding accuracy, Hit@5, and Precision@5 measures, and qualitative examples showed that highly ranked candidates frequently captured the target object concept. Importantly, the reconstruction remained significantly above chance when the candidate search space was expanded to all 1,000 ImageNet categories (Figure S4), indicating that performance was not dependent on restricting the search to shared concepts. These results show that EEG responses measured for one set of images could be used to reconstruct visual content by searching a much larger set of images that had never been presented during the original EEG experiment.

We next texted whether ImageNet-SimEEG could further provide a useful neural supervisory signal for improving the behavioral alignment of artificial vision models. We optimized a pretrained CORnet-S model against the synthetic EEG responses generated for ImageNet images using a representational alignment framework (Lu et al., 2026; Lu & Wang, 2025), while preserving its original image-classification objective (Figure 4C). The resulting ReAlnet-ALL model showed higher model-behavioral similarity on Brain-Score (Schrimpf et al., 2020) than both the baseline CORnet-S solely trained on images, with improvements evident in both the aggregate behavioral score and across individual behavioral benchmarks (Geirhos et al., 2021; Rajalingham et al., 2018). Strikingly, ReAlnet-ALL also outperformed the original ReAlnet models optimized directly using empirical human EEG. This advantage likely reflects the substantially larger and more diverse neural supervision provided by ImageNet-SimEEG, which expands the learned EEG constrains from tens of thousands of empirically measured image-EEG pairs to more than one million visual inputs. Thus, synthetic EEG did not simply substitute for empirical supervision; by scaling empirically learned neural constraints to a much broader stimulus space, it provided a stronger training signal for shaping behaviorally human-like visual representations.

## Discussion

Computational encoding models offer a powerful way to formalize the mapping from sensory inputs to neural responses, but their scientific utility depends on more than predictive accuracy alone (Kriegeskorte & Douglas, 2019; Naselaris et al., 2011). Here, we developed Img2EEG as a participant-specific image-to-EEG framework that combines predictive modeling with controlled intervention and large-scale extrapolation. Img2EEG generalized to images and object concepts not encountered during training while preserving both stimulus-specific and participant-specific response structure. Selective perturbations of internal representations and visual inputs revealed temporally structured dependencies of the predicted EEG response, and individualized models reproduced established visual electrophysiological phenomena, including contralateral responses and the face-sensitive N170. Finally, scaling the learned mappings to ImageNet produced more than 12 million model-derived EEG responses that retained information useful for cross-dataset EEG identification and for improving the model-behavior alignment of an artificial vision model. Together, these findings establish Img2EEG as more than a synthetic EEG generator: it provides an experimentally manipulable and scalable in silico model that connects finite empirical EEG measurements to model-based mechanistic hypothesis testing and large-scale computational exploration of human visual dynamics.

A central consideration for image-to-neural encoding models is what constitutes successful prediction. Visually evoked EEG responses contain substantial structure that is shared across stimuli, such that a model can achieve high correspondence with measured EEG by reproducing common temporal and spatial response patterns without necessarily preserving the information that distinguishes individual visual inputs. Img2EEG therefore was evaluated not only by correspondence within individual spatiotemporal EEG epochs, but also by its ability to reproduce stimulus-dependent variation across images and to support zero-shot identification directly in EEG space. The ability to distinguish among 200 previously unseen images provides a particularly stringent test of the information retained by the generated responses: the predicted EEG associated with the correct stimulus must be more consistent with the measured response than predictions generated for competing stimuli. This distinction between response fidelity and stimulus specificity may be important more generally for evaluating synthetic neural-response models. High waveform similarity alone does not necessarily imply preservation of the neural information that differentiates stimuli, and future generative or predictive neural models may benefit from explicitly testing both properties. The participant-specific analyses further showed that individualized models retained systematic differences between participants, both in their predicted EEG responses and in the representational geometries expressed at different stages of the model. These differences should not be interpreted as identifying the biological origin of inter-individual variability, but they show that participant-specific information is retained rather than collapsed into a single population-average response model. Preserving this variability is important for in silico experimentation because it allows the same stimulus manipulation or model intervention to be evaluated across multiple individualized response systems, rather than only against an average observer. This creates an opportunity to ask not only whether a predicted neural effect emerges, but also how consistency it generalizes across individuals and whether particular effects depend on participant-specific response structure.

The ability to intervene on a trained model provides a complementary step from prediction toward explanation. Because Img2EEG maintains separable hierarchical visual and semantic representations before their integration, image-specific information carried by individual components can be selectively removed while all other model parameters remain fixed. These interventions revealed that different representations contributed to EEG prediction over different, partially overlapping post-stimulus intervals. Complementary manipulations of the input images further allowed the contribution of physically interpretable stimulus properties, including color and spatial-frequency information, to be localized in time. Of course, ablating a representation in silico simply demonstrates that information carried by that representation is necessary for maintaining the model’s prediction under that intervention; it does not demonstrate that the corresponding artificial layer is itself the neural source of the measured EEG effect. Thus, these analyses should be interpreted as identifying dependencies within the trained encoding model, rather than establishing causal mechanisms in the biological visual system, although its value is of course in using these potential links to generate testable hypotheses. These precise in silico perturbations provide a controlled way to generate temporally specific, experimentally testable hypotheses that can be assessed in vivo with fMRI, MEG, intracranial recordings or animal neurophysiology. Critically, having an openly available resource to serve as an in silico test bed for potential research questions carries immensely appeal, potentially allowing researchers to focus costly and time-intensive in vivo experiments on the most informative stimuli, time windows and representational contrasts identified in silico.

As an initial demonstration of this potential, we reported a series of in silico experiments. In addition to reproducing an early contralateral response to lateralized visual stimuli, Img2EEG naturally and veridically reproduced a complex face-sensitive N170 response in posterior electrodes. Note that a model that predicts individual EEG epochs accurately would not necessarily be expected to reproduce the neural signatures that emerge in controlled experimental contrasts, but Img2EEG did so when fed novel face and object images. The N170 results are particularly informative because the effect generalized beyond a binary face-object contrast. Face-pareidolia stimuli produced an intermediate response between real faces and matched non-face objects, suggesting that the simulated effect varies with face-like visual structure rather than simply reproducing a discrete training label. Moreover, a face-sensitive response remained after direct face-concept EEG examples were removed from Img2EEG training, demonstrating that explicit face-specific EEG supervision was not necessary for the learned image-to-EEG mapping to express this effect. Targeted representation ablation further identified a particularly strong dependence of the simulated N170 on the CORnet V4-like representation. While such a finding could stimulate future research on mechanistic accounts of the biological N170, our goal here was focused on providing a proof of principle for how Img2EEG can turn a well-established electrophysiological phenomenon into a tractable in silico test case. By combining controlled stimulus manipulations, training-set interventions and representation ablations, the framework can generate specific hypotheses about which forms of information are sufficient to support an EEG effect within the model and identify candidate manipulations for subsequent empirical testing.

A broader implication of our work – evidenced by the creation of the ImageNet-SimEEG open resource – is that learned image-to-neural mappings can extend the effective stimulus space available for hypothesis generation and experimental design. This is not to say that synthetic responses can replace empirical EEG, but it can offer a valuable complementary tool that may even enhance the effectiveness of empirical studies. Once an image-to-EEG mapping has been learned, it can be evaluated over visual spaces far larger than those that are practical to sample experimentally. Human electrophysiology is necessarily constrained in the number of stimuli and conditions that can be sampled, particularly when repeated measurements are required to obtain reliable responses. For perspective, acquiring neural responses to the 1.28 million images of ImageNet, matching the 80 repetitions used for the THINGS EEG2 test set to get stable ERPs, would require approximately 5,700 hours (1.28 × 10^6^ × 80 × 0.2s = 2.048 × 10^7^ s), or 237 days, of uninterrupted stimulus presentation for a single participant – even before accounting for electrode preparation, breaks, rejected trials or other practical constraints. In this sense, a large model-derived response space such as ImageNet-SimEEG can serve as a computational screening stage, allowing researchers to explore candidate stimuli, representational contrasts and predicted neural effects before committing to empirical data collection. This may be particularly useful for identifying informative regions of a large stimulus space, selecting stimuli expected to maximize neural contrasts, or prioritizing hypotheses for experiments in which acquisition is costly or sampling is limited. The downstream analyses in the present study provide an initial proof of principle that such synthetic response spaces can retain useful structure beyond the dataset from which the encoding model was learned, offering exciting potential in this vein. More broadly, model-derived neural response spaces could support multimodal integration with fMRI, MEG and intracranial electrophysiology, data-driven discovery of temporally resolved representational dimensions, model-guided stimulus selection and in silico pilot testing. However, we emphasize that this utility should be interpreted as complementary to, rather than substitutive of, direct neural measurement. Synthetic EEG necessarily inherits the biases and limitations of both the empirical training data and the pretrained representations used by the model, and empirical recordings remain essential for validating model-derived predictions and identifying where those predictions fail.

Because Img2EEG and the trained participant-specific models are released publicly with a ready-to-use inference pipeline, these applications can be explored directly with new visual stimuli rather than requiring users to retrain the model from scratch. Given an image, its object-concept label and an image caption, the released Img2EEG framework produces predicted responses for all ten participant-specific models, each comprising 17 posterior channels over the first 500 ms after stimulus onset. Researchers can therefore generate individualized EEG predictions for novel stimuli directly, without retraining the encoding models, and can modify or replace the representational components to address new experimental questions.

Several limitations define the scope of the present framework. First, ImageNet-SimEEG and all other generated responses remain model-derived predictions rather than empirical neural measurements, and their validity is therefore bounded by the training data, pretrained representations and learned encoding functions described above.. Second, the present models predict activity from 17 posterior electrodes over the first 500 ms after image onset and therefore capture only a restricted view of whole-brain visual dynamics. Third, the current study relies primarily on ten participants from a single large-scale EEG dataset. Establishing the generality of the framework will require evaluation across independent datasets, acquisition systems, tasks and participant populations. Finally, although model interventions offer substantially greater experimental control than correlational inspection of learned representations, causal conclusions remain confined to the model itself. Their principal neuroscientific value may therefore lie in narrowing the space of hypotheses, identifying informative stimulus manipulations and guiding empirical experiments capable of testing those hypotheses directly.

Img2EEG should nevertheless be viewed as a framework rather than a fixed representational architecture. The present implementation combines CORnet-derived hierarchical visual features with CLIP visual embeddings, object-concept representations and caption-derived semantic embeddings, but none of these specific feature spaces is intrinsic to the general approach. Future versions could incorporate more predictive pretrained representations, representations designed explicitly for interpretability, or additional features motivated by specific hypotheses about visual computation. For example, representations isolating shape, texture, spatial frequency, motion, depth, category structure or other theoretically relevant dimensions could be incorporated and selectively manipulated within the same framework. The model could likewise be extended to denser EEG, MEG, intracranial electrophysiology or multimodal neural measurements, and adapted as new artificial vision and foundation-modal representations become available. This modularity creates an opportunity not only to improve prediction, but also to make future interventions increasingly informative about the representations supporting human visual dynamics.

More broadly, the promise of image-to-neural encoding models is not simply that they can generate larger quantities of synthetic neural data. Their greater potential lies in linking the fidelity of empirical neural measurements with the experimental control and scale of computational models. A useful in silico model should predict responses to new stimuli, preserve the information that differentiates those responses, permit controlled manipulation of the representations on which its predictions depend and extrapolate learned neural constraints beyond the finite stimulus spaces that can be measured directly. Img2EEG provides one step toward this goal for human visual electrophysiology, offering a framework in which prediction, intervention and scaling can be combined to complement direct measurement of the temporal dynamics of human vision.

## Methods

### THINGS EEG2 dataset and EEG preprocessing

#### Participants and experimental paradigm

We used the publicly available THINGS EEG2 dataset (Ref Gifford et al., 2022), which contains EEG recordings from 10 healthy adults (28.5±4.0 years; 8 female and 2 male) performing a rapid serial visual presentation (RSVP) task. Participants viewed naturalistic object images drawn from the THINGS database while maintaining fixation and performing an orthogonal target-detection task to ensure attention to the visual stream. Each image was presented centrally for 100 ms with a stimulus-onset asynchrony of 200 ms (5 Hz). Each participant completed four experimental sessions, yielding extensive repeated measurements of visually evoked EEG responses.

#### Visual stimuli and train-test split

The THINGS EEG2 dataset (Gifford et al., 2022) contains responses to 16,740 image conditions sampled from the THINGS object database (Hebart et al., 2019; Stoinski et al., 2023). We used the original training and test partitions: The training partition comprised 16,540 distinct natural images, with each image presented four times to each participant. The test partition comprised 200 images representing 200 distinct object concepts, with each image presented 80 times per participant. Critically, neither the image identities nor the object concepts represented in the test partition overlapped with those in the training partition, allowing model performance to be evaluated on previously unseen visual inputs and concepts.

For Img2EEG training and evaluation, EEG repetitions corresponding to the same image were averaged to obtain a single response for each image condition. Thus, for each participant, the final dataset consisted of 16,540 image–EEG pairs for model training and 200 independent image–EEG pairs for model testing.

#### EEG acquisition and preprocessing

EEG acquisition and preprocessing followed the procedures described for the original THINGS EEG2 dataset (Gifford et al., 2022). Briefly, EEG was recorded using a 64-channel EASYCAP with a BrainVision actiCHamp amplifier at a sampling rate of 1,000 Hz, referenced online to Fz and filtered during acquisition between 0.1 and 100 Hz. We used the preprocessed EEG data released with the dataset and did not repeat preprocessing from the raw recordings. In the original preprocessing pipeline, trials were baseline-corrected by subtracting the mean voltage during the 100-ms pre-stimulus interval, downsampled to 100 Hz, and restricted to 17 posterior occipital and parietal electrodes (O1, Oz, O2, PO7, PO3, POz, PO4, PO8, P7, P5, P3, P1, Pz, P2, P4, P6 and P8).

For the present study, we retained the post-stimulus interval from 0 to 500 ms. Each image-evoked EEG response was therefore represented as a 17-channel × 50-time-point matrix, corresponding to a temporal resolution of 10 ms per sample. These matrices served as the neural targets for training and evaluating the participant-specific Img2EEG models.

### Img2EEG architecture and training

#### Overview of the Img2EEG architecture

Img2EEG was designed to predict the full spatiotemporal EEG response evoked by a natural image by integrating complementary visual and semantic representations. For each input image, two parallel encoding pathways were constructed: a hierarchical visual encoder based on intermediate representations from a pretrained CORnet-S model (Kubilius et al., 2018, 2019) and a semantic encoder combining CLIP-derived visual features (Radford et al., 2021), object-concept embeddings, and image-description embeddings. The outputs of the two pathways were independently projected to 512-dimensional representations, concatenated, and passed through three fully connected integration layers. The final layer produced an 850-dimensional output corresponding to the predicted EEG response across 17 channels and 50 post-stimulus time points.

#### Hierarchical visual encoder

The hierarchical visual encoder was based on a pretrained CORnet-S model (Kubilius et al., 2018, 2019), a recurrent convolutional neural network organized into different hierarchical stages corresponding approximately to brain areas V1, V2, V4 and inferotemporal cortex. For Img2EEG, we extracted activations from the V1, V2 and V4 stages to obtain low-to mid-level visual information. For a 224 × 224 RGB input image, these stages produced feature maps of 64 × 56 × 56, 128 × 28 × 28 and 256 × 14 × 14 units, respectively. The three feature maps were flattened and concatenated, yielding a 351,232-dimensional hierarchical visual representation, which was projected through a fully connected layer followed by a rectified linear unit (ReLU) to obtain a 512-dimensional visual feature vector.

CORnet-S weights were initialized from the publicly released pretrained CORnet-S model and remained frozen throughout Img2EEG training; only the subsequent projection and integration layers were optimized against EEG responses. Images entering CORnet-S were resized to 224 × 224 pixels and normalized using the ImageNet channel means and standard deviations.

#### Semantic encoder

The semantic encoder combined three complementary representations of each image. First, a 512-dimensional visual representation was extracted using the image encoder of pretrained CLIP ViT-B/32 (Radford et al., 2021). The CLIP parameters were frozen during Img2EEG training. Images entering CLIP were resized to 224 × 224 pixels and normalized using the preprocessing parameters associated with the pretrained CLIP model.

Second, each image was associated with its THINGS object concept and represented using a 300-dimensional GloVe embedding from the 840B-token, 300-dimensional GloVe model (Pennington et al., 2014). Object labels were looked up as complete tokens; labels that were not present in the GloVe vocabulary were represented using the model’s “unk” vector.

Third, a natural-language description was generated independently for each image using BLIP-2 with the OPT-2.7B language model (J. Li et al., 2023). Each generated caption was subsequently encoded using the pretrained MPNet sentence-embedding model (all-mpnet-base-v2) (K. Song et al., 2020), yielding a 768-dimensional image-description representation.

The 512-dimensional CLIP visual representation, 300-dimensional GloVe object-concept representation, and 768-dimensional MPNet image-description representation were concatenated into a 1,580-dimensional semantic feature vector. This vector was projected through a trainable fully connected layer followed by a rectified linear unit (ReLU) to obtain a 512-dimensional semantic representation.

#### Feature integration and EEG readout

The 512-dimensional hierarchical visual representation and 512-dimensional semantic representation were concatenated into a joint 1,024-dimensional feature vector. This combined representation was passed through two successive 512-unit fully connected layers, each followed by ReLU, and then through a final linear readout layer with 850 output units. The 850-dimensional output was interpreted as the complete predicted EEG epoch, corresponding to 17 posterior EEG channels × 50 time points spanning the first 500 ms following image onset.

This architecture therefore preserved distinct visual and semantic encoding pathways before integrating them into a common representation used to predict the temporally resolved EEG response. Importantly, the visual and semantic feature extractors were held fixed, whereas the projection layers in the hierarchical visual and semantic encoders and all subsequent integration layers were optimized separately for each participant.

#### Participant-specific model training

A separate Img2EEG model was trained for each of the ten THINGS EEG2 participants. Before training, EEG responses were standardized separately for each participant using the global mean and standard deviation calculated across that participant’s complete training set. The same training-derived normalization parameters were subsequently applied to the corresponding test responses.

Trainable Img2EEG parameters were optimized using the Adam optimizer (Kingma & Ba, 2014) to minimize mean-squared error between the predicted and real 850-dimensional EEG responses. The provided training implementation used a batch size of 16 and a learning rate of 10^-4^, with training samples shuffled independently within each epoch. Random seeds were controlled independently for each participant to improve reproducibility.

#### Public implementation and inference

We provide a public implementation of Img2EEG together with the ten trained participant-specific models. For inference on a new stimulus, users provide an input image, its corresponding object-concept label and an image caption. These inputs are processed through the pretrained visual and semantic feature extractors described above and passed to each of the ten individualized Img2EEG models, yielding ten participant-specific simulated EEG responses. Each output consists of 17 posterior channels × 50 time points spanning 0-500 ms after stimulus onset. Thus, the released implementation allows EEG responses for novel visual stimuli to be generated without retraining the encoding models.

### Evaluation of EEG prediction and stimulus specificity

#### Signal-pattern similarity

To quantify the overall correspondence between generated and empirical EEG responses for individual images, we calculated a signal-pattern similarity measure. For each test image, the generated EEG response and the corresponding empirical EEG response were each reshaped from a 17-channel × 50-time-point matrix into an 850-element vector. Spearman’s rank correlation was then calculated between the two vectors. This yielded one correlation coefficient for each of the 200 held-out test images and each participant. Signal-pattern similarity therefore quantified how well Img2EEG reproduced the overall spatiotemporal organization of the EEG response evoked by a given image.

#### Stimulus-profile similarity

To quantify how well Img2EEG preserved stimulus-dependent variation in EEG responses, we calculated a complementary stimulus-profile similarity measure. For each channel and time point, the generated response values across the 200 held-out test images were arranged into a 200-element vector and compared with the corresponding 200-element vector from the real EEG responses using Spearman’s rank correlation. This procedure yielded one correlation coefficient for each of the 17 channel × 50 time-point combinations. To obtain a global summary of stimulus-dependent correspondence across the recorded spatiotemporal response, correlation coefficients were averaged across all channels and time points to obtain a participant-level stimulus-profile similarity score.

Additionally, to determine whether stimulus-profile similarity depended on the correct correspondence between stimulus identity and EEG response, we performed a stimulus-shuffling control separately for each participant. Image identities were randomly permuted across the 200 test stimuli 1,000 times while preserving the generated and empirical response values themselves. Stimulus-profile similarity was recomputed for each permutation using the same procedure described above, and the 1,000 shuffled similarity values were averaged to obtain a single participant-specific shuffled baseline. The intact stimulus-matched similarity was then compared with this shuffled baseline across participants using a paired two-sided *t*-test.

Whereas signal-pattern similarity measures correspondence across the full spatiotemporal EEG pattern within individual stimuli, stimulus-profile similarity measures correspondence in the relative pattern of responses across stimuli at each channel and time point. The latter therefore provides a more direct assessment of whether Img2EEG captures image-dependent variation rather than only response structure that is shared across stimuli.

#### Participant-specificity analysis

To assess whether Img2EEG captured participant-specific neural response structure, we compared correspondence between real EEG responses and responses generated by matched and unmatched participant-specific models. For each empirical participant, signal-pattern and stimulus-profile similarity were calculated using EEG generated by that participant’s own Img2EEG model and separately using EEG generated by each of the other nine Img2EEG models. Matched-model similarity was compared with the average similarity obtained from the unmatched models. This analysis was performed separately for signal-pattern and stimulus-profile similarity, providing complementary tests of whether individualized Img2EEG models captured participant-specific spatiotemporal response patterns and participant-specific stimulus-dependent response structure.

#### Component ablation for prediction

To evaluate the contribution of individual visual and semantic representations to EEG prediction, we selectively ablated each representational component of Img2EEG while leaving all remaining model components and parameters unchanged. For a given component, image-specific information was removed by replacing the feature representation for each test image with the mean representation calculated across the training set. The ablated representation was then propagated through the remaining trained model to generate a perturbed EEG response.

Prediction performance of each ablated model was evaluated using the same signal-pattern and stimulus-profile similarity measures described above. The effect of each ablation was quantified by comparing performance of the ablated model with that of the intact Img2EEG model. This analysis was conducted separately for representations derived from CORnet V1, V2 and V4, CLIP visual features, GloVe-based object-concept features and MPNet-based image-description features.

#### EEG-to-image zero-shot retrieval

To test whether generated EEG responses contained sufficient stimulus-specific information to identify previously unseen images, we performed EEG-to-image zero-shot retrieval on the 200-image THINGS EEG2 test set. For each participant and each empirical test EEG response, Img2EEG was used to generate candidate EEG responses for all 200 test images. The empirical EEG response and each candidate generated response were represented in the same 850-dimensional channel × time space, and their dissimilarity was quantified using mean-squared error (MSE). Candidate images were ranked from the smallest to the largest MSE, with the image whose generated EEG response most closely matched the empirical response assigned the highest rank.

Top-1 retrieval was scored as correct when the image corresponding to the empirical EEG response was ranked first among all 200 candidates. Top-5 retrieval was scored as correct when the target image appeared among the five highest-ranked candidates. Because neither the test images nor their object concepts were included during Img2EEG training, this analysis provides a direct test of whether the model preserved discriminative EEG information for unseen visual inputs.

#### Comparison with existing EEG-to-image decoding models

To contextualize the zero-shot identification performance of Img2EEG, we compared its Top-1 and Top-5 accuracy with four previously reported EEG-based visual decoding approaches evaluated on the THINGS EEG2 dataset: BraVL (Du et al., 2023), NICE (Y. Song et al., 2024), ATM (D. Li et al., 2024) and CognitionCapturer (Zhang et al., 2025). All comparisons used subject-specific results for the 200 held-out THINGS EEG2 concepts, which are disjoint from the 1,654 concepts used for model training. Because the published methods differ in their decoding objectives and EEG preprocessing pipelines, these comparisons were used as benchmarks against the best reported performance under each method rather than as controlled architectural comparisons.

BraVL learns a joint brain–visual–linguistic latent representation and performs zero-shot category decoding using visual and textual information from novel concepts. On THINGS EEG2, BraVL reported mean 200-way Top-1 and Top-5 accuracies of 5.82% and 17.45%, respectively. NICE aligns EEG and visual representations through contrastive learning and performs zero-shot recognition by matching EEG embeddings to representations of unseen visual concepts; its original subject-dependent THINGS EEG2 evaluation reported mean Top-1 and Top-5 accuracies of 13.8% and 39.5%, respectively. ATM extends this contrastive-learning framework with a channel-wise attention and temporal–spatial EEG encoder and explicitly evaluates image retrieval by matching EEG embeddings against CLIP embeddings of the 200 test images. We therefore used its subject-dependent retrieval results (Top-1, 28.64%; Top-5, 58.47%) as the ATM benchmark. CognitionCapturer uses modality-specific EEG encoders to align EEG with image, text and depth representations. For comparison with our visually driven identification task, we used the image-modality expert results rather than the multimodal upper-bound score, yielding mean 200-way Top-1 and Top-5 accuracies of 33.30% and 60.58%, respectively.

The Img2EEG analysis differs conceptually from these decoding approaches in that no EEG-to-image decoder or shared EEG-image embedding space was trained. Instead, Img2EEG generates a candidate EEG response independently for each unseen image, and stimulus identity is inferred directly by comparing these generated responses with the measured EEG response. Accordingly, the benchmark comparison assesses whether the image-specific neural predictions generated by Img2EEG contain sufficient information to support visual identification at a level comparable to or exceeding dedicated EEG decoding models.

#### Component ablation for zero-shot retrieval

To determine which Img2EEG representations contributed to stimulus identification, we repeated the zero-shot retrieval analysis after selectively ablating each visual or semantic component using the same mean-replacement procedure described above. For each ablated model, candidate EEG responses were regenerated for all 200 test images and ranked against each empirical EEG response using MSE. Top-1 and Top-5 retrieval performance were then recalculated and compared with performance of the intact Img2EEG model. This analysis allowed us to assess which representational components were most important for preserving stimulus-discriminative information in the generated EEG responses.

### Representational analysis of individual differences

#### Feature extraction across individualized models

To examine how individual differences were expressed across different stages of Img2EEG, we presented the 200 held-out THINGS EEG2 test images to each of the ten participant-specific models and extracted representations from three stages of the architecture: the output of the hierarchical visual encoder, the output of the semantic encoder, and the output of the second integration layer. Feature extraction was performed independently for each participant-specific Img2EEG using the same set of 200 test images.

#### Representational dissimilarity matrices

For each participant and representational stage, we constructed a 200 × 200 representational dissimilarity matrix (RDM) (Kriegeskorte et al., 2008) describing the pairwise representational relationships among the 200 held-out images. Each matrix element reflected the dissimilarity between the feature representations elicited by a pair of images, and diagonal elements were excluded from subsequent analyses. This yielded one RDM for each participant at each of the three representational stages.

#### Inter-subject representational similarity

To quantify the extent to which representational geometry was shared across individualized Img2EEG models, we compared RDMs between participants separately for each representational stage. The upper-triangular elements of each pair of RDMs were vectorized, and their similarity was quantified using Spearman’s rank correlation. With ten individualized models, this yielded 45 pairwise inter-subject similarity values for each stage. Higher correlations indicated more similar organization of the 200-image representational space across participants, whereas lower correlations indicated greater inter-individual variability.

Inter-subject representational similarity was then compared across the hierarchical visual, semantic and integration representations to characterize how shared versus individualized representational structure varied across the Img2EEG architecture.

### Temporally resolved feature ablation analyses

#### Internal feature ablation

To determine when information carried by individual Img2EEG representations contributed to EEG prediction, we performed feature-specific ablations within the hierarchical visual and semantic encoders. For each target representation, image-specific information was removed while preserving the overall distributional scale of that feature space. Specifically, the representation elicited by each test image was replaced with the mean representation calculated across all images in the THINGS EEG2 training set. All remaining representations and trained model parameters were left unchanged, and the modified feature set was propagated through the downstream Img2EEG layers to generate an ablated EEG response.

Ablations were performed separately for representations derived from CORnet V1, V2 and V4, the CLIP visual embedding, the GloVe-based object-concept representation, and the MPNet-based image-description representation. Because the replacement vector was constant across test images, this manipulation selectively removed stimulus-dependent information carried by the targeted representation without requiring retraining of the model. The resulting EEG predictions were compared with those of the intact Img2EEG model to determine when removal of each representation reduced correspondence with empirical EEG.

#### Input-level feature ablation

We complemented the internal representation ablations with manipulations applied directly to the visual inputs. For each of the 200 held-out test images, we generated modified versions in which color, low-spatial-frequency information, or high-spatial-frequency information was selectively removed. Each manipulated image was then passed through the intact participant-specific Img2EEG model, with all model parameters held fixed, to generate a corresponding EEG response.

Color information was removed by converting the original images to grayscale. Spatial-frequency manipulations were performed independently for each image channel in the two-dimensional Fourier domain. Images were first transformed using a discrete Fourier transform and the zero-frequency component was shifted to the center of the spectrum. Circular ideal frequency masks were then applied to selectively remove low- or high-spatial-frequency information. For the low-spatial-frequency ablation, frequencies at or below approximately 7 cycles per image were removed using a high-pass mask. For the high-spatial-frequency ablation, frequencies above approximately 17.5 cycles per image were removed using a low-pass mask. The filtered spectra were transformed back into the image domain using an inverse discrete Fourier transform, after which each image was normalized to the 0–255 intensity range and converted to 8-bit format.

EEG responses generated from each manipulated image were compared with the corresponding empirical EEG responses using the same time-resolved prediction-similarity analysis as for the internal feature ablations. In this way, internal representation ablations and direct stimulus manipulations provided complementary tests of when model-derived representations and physically interpretable visual features contributed to accurate EEG prediction.

#### Time-resolved prediction similarity

To quantify the effect of each ablation over time, we calculated generated-to-empirical EEG correspondence independently at each post-stimulus time point. For a given participant and time point, EEG responses across all 200 test images and 17 EEG channels were arranged into a 200 × 17 matrix for both the generated and empirical data. Each matrix was flattened into a 3,400-element vector, and Spearman’s rank correlation was calculated between the generated and empirical vectors.

This procedure was applied separately to EEG responses generated by the intact model and by each internal- or input-level ablation condition. The time-resolved ablation effect was quantified as the difference in prediction similarity between the intact and ablated conditions:

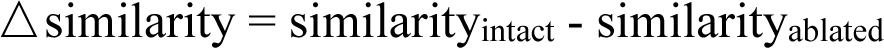

Positive values therefore indicated that removal of the targeted feature or representation reduced the model’s ability to account for empirical EEG responses at that time point. Repeating this analysis across the 50 post-stimulus time points yielded a temporal contribution profile for each ablated representation or stimulus feature and each participant. Statistical inference on these time-resolved effects is described in Statistics and reproducibility.

### In silico experiments on canonical visual EEG phenomena

#### Contralateral visual response experiment

To test whether Img2EEG reproduced a basic spatially organized property of visually evoked EEG, we conducted an in silico experiment using laterally presented object stimuli. Forty object images were each resized to 300 × 300 pixels and placed on a white 1,200 × 1,200-pixel canvas. For each object, two stimulus versions were generated by positioning the object center 300 pixels to the left or right of the center of the canvas while maintaining a constant vertical position, yielding 80 lateralized stimuli in total.

Each stimulus was presented to all participant-specific Img2EEG models to generate simulated EEG responses. For each stimulus position, non-midline electrodes were relabeled as contralateral or ipsilateral according to their hemisphere relative to the visual-field location of the object. Simulated contralateral and ipsilateral ERPs were then averaged across the 14 non-midline posterior electrodes, and the contralateral effect was quantified from the difference between the two response time courses.

#### Face-selective N170 experiment

To test whether Img2EEG recapitulated a canonical category-sensitive visual ERP, we conducted an in silico face-selective N170 experiment using 40 grayscale face images and 40 grayscale object images from a stimulus set used in fMRI functional localizer experiments in our laboratory (Dube et al., 2022; Finlayson et al., 2017; Nag et al., 2019). These stimuli were independent of the THINGS EEG2 training images. Each image was presented to all ten participant-specific Img2EEG models, and the resulting simulated EEG responses were averaged separately across face and object stimuli to obtain condition-specific ERPs.

The temporal N170 effect was assessed from the difference between the face- and object-evoked ERP time courses. We focused on the 170-200 ms post-stimulus interval, corresponding to the canonical N170 time window reported in previous face-processing studies (Hinojosa et al., 2015; Mercure et al., 2011; Rossion & Jacques, 2012). Electrodes P8 and PO8 were selected as the primary channels because the N170 is typically maximal over lateral posterior occipitotemporal electrodes, with strong face-selective responses commonly observed at right posterior sites. To characterize the spatial distribution of the simulated N170 response, we additionally quantified face selectivity separately at each EEG channel. For this analysis, ERP amplitudes were averaged over the 170–200 ms post-stimulus interval for face and object stimuli separately, and the condition difference was calculated for each channel. This yielded a channel-wise measure of the simulated N170 effect across the posterior electrode array.

#### Representation ablation of the N170 effect

To determine which Img2EEG representations contributed to the emergence of the simulated N170 effect, we repeated the face-versus-object experiment after selectively ablating individual representational components. Internal feature ablations were performed using the same mean-replacement procedure described in the temporally resolved feature-ablation analyses. For each targeted component, the image-specific feature representation was replaced with the corresponding mean representation computed across the THINGS EEG2 training set, while all other representations and trained model parameters were held fixed.

Ablations were performed separately for representations derived from CORnet V1, V2 and V4, the CLIP visual representation, the GloVe-based object-concept representation, and the MPNet-based image-description representation. For each ablated model, face- and object-evoked EEG responses were regenerated, and the N170 effect was recalculated at electrode P8 as the mean face-object amplitude difference over the 170-200 ms interval. The effect of each representation ablation was quantified relative to the corresponding N170 effect generated by the intact Img2EEG model.

#### Pareidolia experiment

To test whether the face-sensitive response generated by Img2EEG generalized to stimuli with graded face-like appearance, we conducted an additional in silico experiment using the face-pareidolia stimulus set (Wardle et al., 2020). The stimulus set comprised 96 color photographs: 32 human faces, 32 inanimate objects containing illusory faces, and 32 matched inanimate objects without illusory faces. The non-face objects were yoked to the pareidolia images such that each illusory-face stimulus was paired with an object of the same identity that was selected to be as visually similar as possible while lacking the perception of a face. All 96 images were presented to each participant-specific Img2EEG model. Simulated ERPs were averaged separately for the face, pareidolia and matched-object conditions.

#### Non-face-trained Img2EEG

To test whether direct exposure to face-evoked EEG examples during Img2EEG training was necessary for the emergence of the simulated N170 effect, we trained an additional set of ten participant-specific models after removing the face concept from the THINGS EEG2 training set. THINGS EEG2 is organized at the object-concept level, with natural images assigned to individual object concepts; removing the face concept therefore excluded the full set of face-category training images and their corresponding EEG responses while leaving all remaining object concepts unchanged. The model architecture, feature extraction procedures and optimization settings were otherwise identical to those used for the original Img2EEG models.

The resulting non-face-trained Img2EEG models were evaluated using the same independent grayscale face and object stimulus sets used in the primary N170 experiment. Simulated ERPs were generated separately for the two conditions, and the temporal face–object difference was evaluated at P8. The channel-wise N170 distribution was additionally quantified from mean face–object amplitude differences over the 170-200 ms interval. These analyses tested whether a face-selective N170-like response could be expressed by the learned image-to-EEG mapping in the absence of direct face-concept EEG supervision during model training.

### ImageNet-SimEEG generation and downstream analyses

#### ImageNet image set

To construct a large-scale synthetic EEG resource, we used the ImageNet 2012 image set (Deng et al., 2009), including both the training and validation partitions. Together, these partitions comprise approximately 1.28 million natural images spanning 1,000 object categories. All images were processed using the same image-preprocessing pipelines used for Img2EEG inference and were independently presented to each of the ten participant-specific Img2EEG models.

#### Generation of ImageNet-SimEEG

For each ImageNet image, we generated a predicted EEG response from each of the ten individualized Img2EEG models. Each prediction consisted of 17 posterior EEG channels × 50 time points spanning 0-500 ms after stimulus onset. Applying the ten participant-specific models to the complete ImageNet image set yielded more than 12 million model-derived EEG epochs in total. We refer to this large-scale synthetic neural response space as ImageNet-SimEEG.

#### Database-based image reconstruction from EEG

To test whether ImageNet-SimEEG could support reconstruction of visual content from independently measured EEG, we used empirical EEG responses from the THINGS EEG2 test set as queries against the synthetic ImageNet EEG response space. Quantitative evaluation was initially restricted to the 57 object concepts shared between THINGS EEG2 and ImageNet, and only ImageNet images belonging to these 57 concepts were included in the initial candidate pool.

For each empirical THINGS EEG2 response, the corresponding participant-specific ImageNet-SimEEG responses were compared with the empirical EEG response in the same neural response space. Candidate ImageNet images were ranked according to EEG similarity, from the most to the least similar response. Because multiple ImageNet images belonged to each of the 57 candidate concepts, reconstruction performance was evaluated at the concept level using three complementary metrics.

Decoding accuracy was defined as the proportion of THINGS EEG2 queries for which the highest-ranked ImageNet image belonged to the same object concept as the query stimulus. Hit@5 was defined as the proportion of queries for which at least one of the five highest-ranked ImageNet images belonged to the correct object concept. Precision@5 quantified the proportion of the five highest-ranked images that belonged to the correct concept, calculated for each query as the number of correctly matched images among the top five divided by five. Metrics were first calculated separately for each participant and then summarized across participants.

These complementary measures capture different aspects of cross-dataset retrieval: decoding accuracy assesses whether the single best-matching synthetic EEG response identifies the correct concept, Hit@5 assesses whether the correct concept is represented among the highest-ranked candidates, and Precision@5 measures how consistently images from the target concept are concentrated near the top of the ranked retrieval set.

As a more stringent reconstruction analysis, we repeated the analysis without constraining the candidate image pool to the 57 overlapping concept categories. Each empirical EEG response from THINGS EEG2 was searched against the full ImageNet-SimEEG validation set spanning all 1,000 ImageNet categories. The same ranking and evaluation metrics were applied.

### EEG-guided optimization of CORnet

To test whether ImageNet-SimEEG could provide a scalable neural supervisory signal for artificial vision models, we optimized pretrained CORnet-S models jointly for ImageNet object recognition and alignment with the synthetic EEG responses. The optimization followed the architecture of the ReAlnet framework (Lu et al., 2026; Lu & Wang, 2025), with ImageNet-SimEEG replacing empirically recorded brain signals as the neural target.

For EEG-guided optimization, we used the first 200 ms of each ImageNet-SimEEG response, corresponding to 17 posterior EEG channels × 20 time points (340 values). For each ImageNet image, activations were extracted from layer V1, V2, V4 and IT of CORnet-S. The corresponding feature maps were flattened and independently projected through trainable fully connected layers to 128-dimensional representations. The four projected representations were concatenated into a 512-dimensional feature vector and passed through a final linear readout to predict the corresponding synthetic EEG target.

CORnet-S and the EEG readout were optimized jointly using a combined objective comprising the original ImageNet object-classification loss and an EEG-alignment loss. Object classification was optimized using cross-entropy loss, whereas correspondence between the predicted and ImageNet-SimEEG responses was optimized using mean-squared error. Thus, model optimization was constrained simultaneously by the original visual-recognition objective and by the participant-specific neural response structure derived from Img2EEG. Models were initialized from publicly available pretrained CORnet-S weights. All training parameters and training process remained consistent with the original neural data-based ReAlnet paper (Lu et al., 2026).

#### Evaluation of model-brain alignment

Model–brain alignment was evaluated using the standardized Brain-Score Vision benchmarking platform (Schrimpf et al., 2020). We used two behavioral benchmarks from the Brain-Score: “Rajalingham2018public-i2n”7 task uses grayscale images where objects are manipulated by varying position, size, viewing angle, and background (Rajalingham et al., 2018), while “Geirhos2021-error_consistency”42 task employs out-of-distribution colorful natural images (Geirhos et al., 2021). Both tasks calculate behavioral similarity between the model and human (and primates) observers using the error consistency method, which measure whether there is above-chance overlap in the specific images that humans and models classify incorrectly.

All models were evaluated using the same Brain-Score pipeline, allowing the ImageNet-SimEEG-optimized models to be directly compared with real EEG-optimized ReAlnets and their pretrained CORnet-S baseline. Brain-Score evaluation was conducted after model training and was not used for parameter optimization or model selection.

### Statistics and reproducibility

All statistical analyses were conducted at the participant level unless otherwise specified. The primary analyses included all ten participants in the publicly available THINGS EEG2 dataset, and no statistical method was used to predetermine sample size. Unless otherwise stated, statistical tests were two-sided and summary values are reported as mean ± SEM across participants. Multiple comparisons were controlled using false discovery rate (FDR) correction where indicated.

#### EEG prediction and participant- and stimulus-specificity

Signal-pattern and stimulus-profile similarities were quantified using Spearman’s rank correlation as described above. To assess participant specificity, for each empirical participant we compared similarity obtained using the matched participant-specific Img2EEG model with similarity obtained from the nine unmatched models. The unmatched value for each participant was defined as the average similarity across the nine models trained on the other participants. Matched and unmatched similarities were compared across participants using paired two-sided t-tests, separately for signal-pattern and stimulus-profile similarity.

For component-ablation analyses of EEG prediction, performance obtained after ablating each visual or semantic representation was compared with that of the intact Img2EEG model using paired two-sided t-tests across participants. Analyses were conducted separately for signal-pattern and stimulus-profile similarity. *P*-values across the six representational ablations were corrected for multiple comparisons using FDR.

#### Zero-shot visual identification

Top-1 and Top-5 zero-shot identification accuracies were calculated independently for each participant. For the 200-way THINGS EEG2 test set, theoretical chance performance was 0.5% (1/200) for Top-1 identification and 2.5% for Top-5 identification. Top-1 and Top-5 Img2EEG accuracies were compared with theoretical chance levels using one-sample two-sided t-tests. Participant-wise Img2EEG accuracies were compared with those of existing EEG-to-image decoding models using paired two-sided t-tests.

For component-ablation analyses of zero-shot identification, Top-1 and Top-5 performance obtained from each ablated Img2EEG model was compared with the corresponding intact-model performance using paired two-sided t-tests across participants. *P*-values across the six ablated representations were FDR-corrected separately for Top-1 and Top-5 analyses.

#### Representational analysis of individual differences

To compare the degree of shared representational geometry across stages of Img2EEG while retaining the participant as the statistical unit, we used a leave-one-participant-out representational similarity procedure. For each participant and representational stage, the participant-specific RDM was compared with a reference RDM obtained by averaging the corresponding RDMs from the remaining nine participants. Similarity between the participant-specific and leave-one-participant-out reference RDMs was quantified using Spearman’s rank correlation between their vectorized upper-triangular elements. This procedure yielded one inter-subject representational similarity value per participant for each of the hierarchical visual, semantic and integration representations.

Representational similarity was compared between stages using paired two-sided t-tests across the ten participants. The three pairwise comparisons among hierarchical visual, semantic and integration representations were corrected for multiple comparisons using FDR. Pairwise participant-by-participant RDM correlations were used for visualization only and were not treated as statistically independent observations.

#### Temporally resolved feature-ablation analyses

For each internal- and input-level ablation, the time-resolved ablation effect was calculated independently for each participant as the difference in generated-to-empirical EEG similarity between the intact and ablated conditions. Positive values therefore indicated a reduction in EEG prediction performance following removal of the targeted representation or stimulus feature.

Statistical significance across time was assessed using cluster-based permutation testing. At each time point, intact and ablated conditions were first compared across participants using a paired two-sided t-test. Temporally adjacent samples reaching an uncorrected threshold of *p* < 0.05 were grouped into clusters. For each cluster, the cluster statistic was calculated as the sum of the constituent t values. Cluster-level significance was assessed using 1,000 permutations of the paired condition labels across participants. For each permutation, the paired labels were randomly exchanged within participants, the complete time-resolved statistical analysis was recomputed, and the maximum cluster statistic was retained to construct the null distribution.

Observed clusters were evaluated relative to this permutation-derived distribution, thereby controlling the family-wise error rate across time. The same procedure was applied independently to each internal representation ablation and each input-level stimulus manipulation.

#### In silico visual EEG experiments

Temporal differences between contralateral and ipsilateral simulated ERPs were assessed using the same cluster-based permutation procedure described above. At each post-stimulus time point, contralateral and ipsilateral responses were compared using paired two-sided t-tests across participant-specific Img2EEG models. Adjacent samples with uncorrected *p* < 0.05 were grouped into clusters, cluster statistics were calculated as the sum of t values, and cluster-level significance was determined from 1,000 permutations of the paired condition labels using the maximum-cluster statistic.

The temporal face–object difference at electrode P8 was evaluated using the same cluster-based permutation procedure. Channel-wise N170 selectivity was quantified as the mean face–object amplitude difference over the 170–200 ms post-stimulus interval.

For the representation-ablation analysis of the N170 effect, the N170 amplitude obtained from each ablated model was compared with that obtained from the intact Img2EEG model using paired two-sided t-tests across participants. *P*-values across the six ablated representations were corrected using FDR.

The non-face-trained Img2EEG models were evaluated using the same statistical procedures as the primary face-selective N170 analysis.

#### ImageNet-SimEEG downstream analyses

For cross-dataset retrieval, concept-level decoding accuracy, Hit@5 and Precision@5 were calculated independently for each participant using the 57 object concepts shared between THINGS EEG2 and ImageNet. Each metric was compared against its corresponding chance level across participants using one-sample (ReAlnet vs. CORnet and ReAlnet-ALL vs. CORnet) or paired (ReAlnet-ALL vs. ReAlnet) two-sided t-tests. Multiple comparisons across the three retrieval metrics were corrected using FDR.

For model-behavior alignment, Brain-Score values obtained for ReAlnet-ALL were compared with those of the real EEG-optimized ReAlnet and the original pretrained CORnet-S separately for the scores from two behavioral benchmarks and also the averaged score. Statistical comparisons were performed using one-sample two-sided t-test.

### Data and code availability

The EEG data (THINGS EEG2 dataset) used are available as open data via the Open Science Framework (OSF) repository: https://osf.io/3jk45 (Gifford et al., 2022). Code for Img2EEG training, model intervention and inference is publicly available at https://github.com/ZitongLu1996/Img2EEG. The same repository also provides access to the ten trained participant-specific Img2EEG model weights and the ImageNet-SimEEG resource, together with the inference pipeline used to generate stimulated EEG responses for user-provided visual stimuli. The pipeline requires an image, an object-concept label and an image caption as input and returns ten individualized 17-channel × 50-time-point EEG responses spanning 0-500 ms.

## Acknowledgement

This work was supported by grants from the National Institutes of Health (R01-EY025648) to Julie D. Golomb. We thank the Ohio Supercomputer Center for providing the essential computing resources and support.

## Supplementary

**Figure S1.**
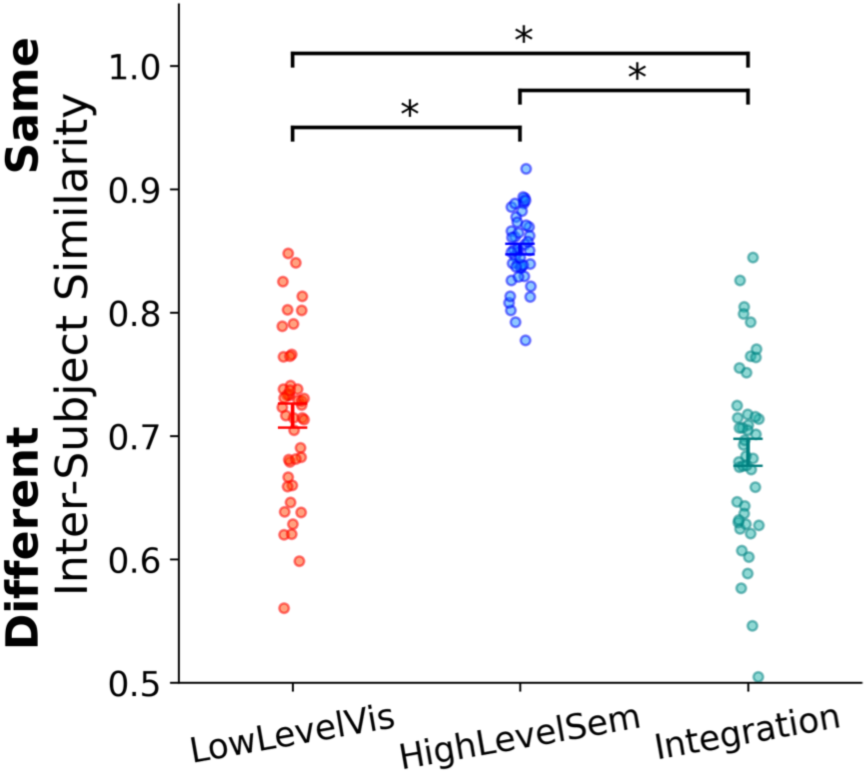
Inter-subject representational similarity across stages of individualized Img2EEG models. Inter-subject representational similarity was compared across the hierarchical visual encoder, semantic encoder and integration layer. Semantic representations showed the highest cross-participant similarity, whereas hierarchical visual and integration representations were more variable across individualized models. Points indicate pairwise participant comparisons; error bars indicate mean ± SEM. Asterisks denote significant differences between representational stages.

**Figure S2.**
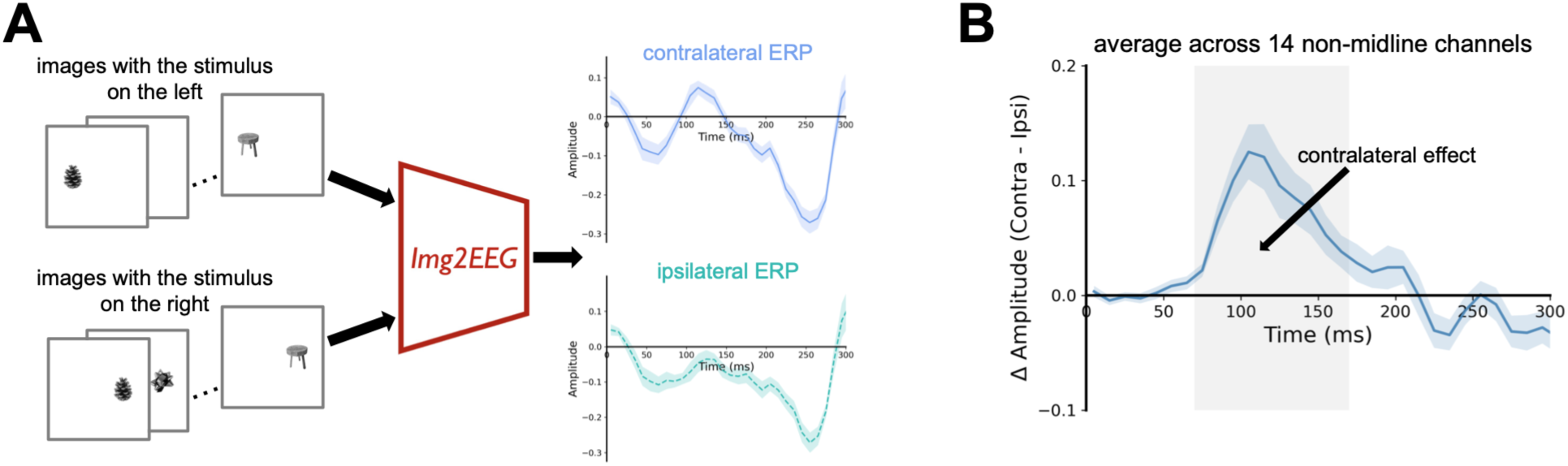
Img2EEG reproduces an early lateralized posterior visual response. (A) Schematic of the in silico experiment used to test the contralateral visual response. Images containing objects represented in the left or right visual field were entered into each participant-specific Img2EEG model, and the resulting simulated EEG responses were organized according to whether each channel was contralateral or ipsilateral to the stimulus location. (B) Time course of the simulated contralateral effect, quantified as the difference between contralateral and ipsilateral responses and averaged across the 14 non-midline channels. Img2EEG reproduced a transient contralateral response following stimulus onset.

**Figure S3.**
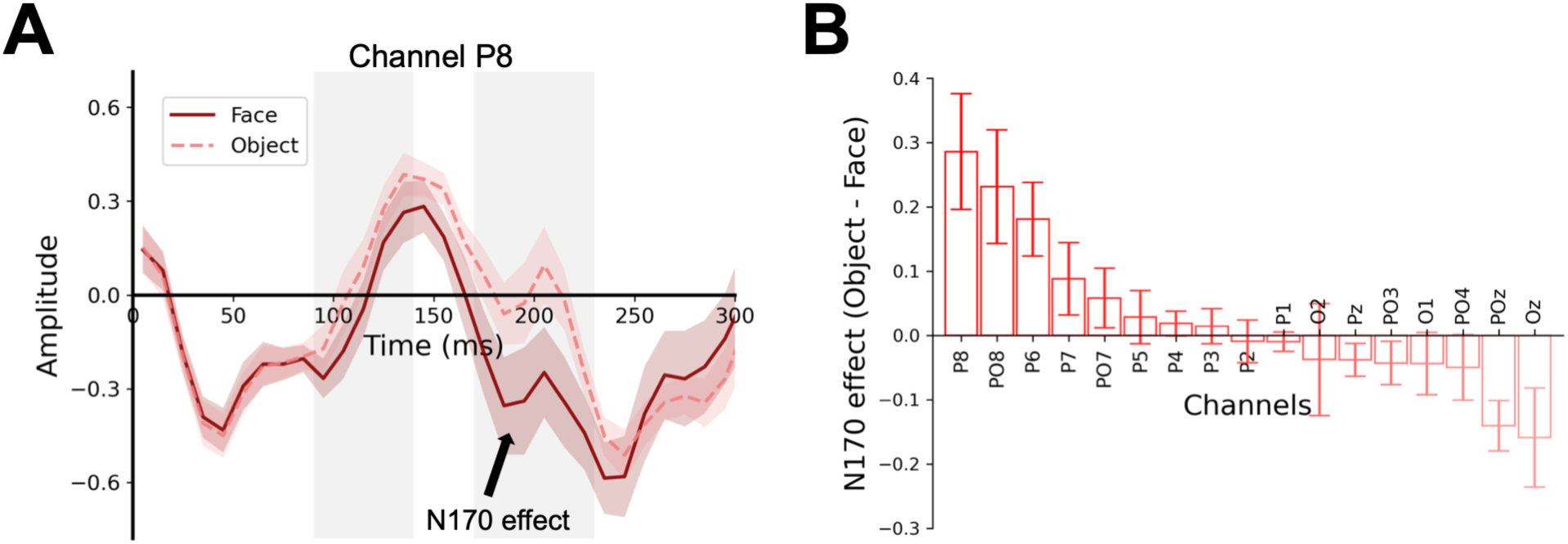
Face-selective N170 responses in Img2EEG models trained without face-concept EEG examples. (A) Simulated face- and object-evoked ERPs at electrode P8 from Img2EEG models retrained after removal of the face concept from the EEG training set. The models continued to show a face-selective N170-like response. (B) Channel-wise N170 effect, quantified as the mean object–face amplitude difference over 170-200 ms, showing the posterior distribution of the effect. Shaded regions and error bars indicate SEM across participant-specific models.

**Figure S4.**
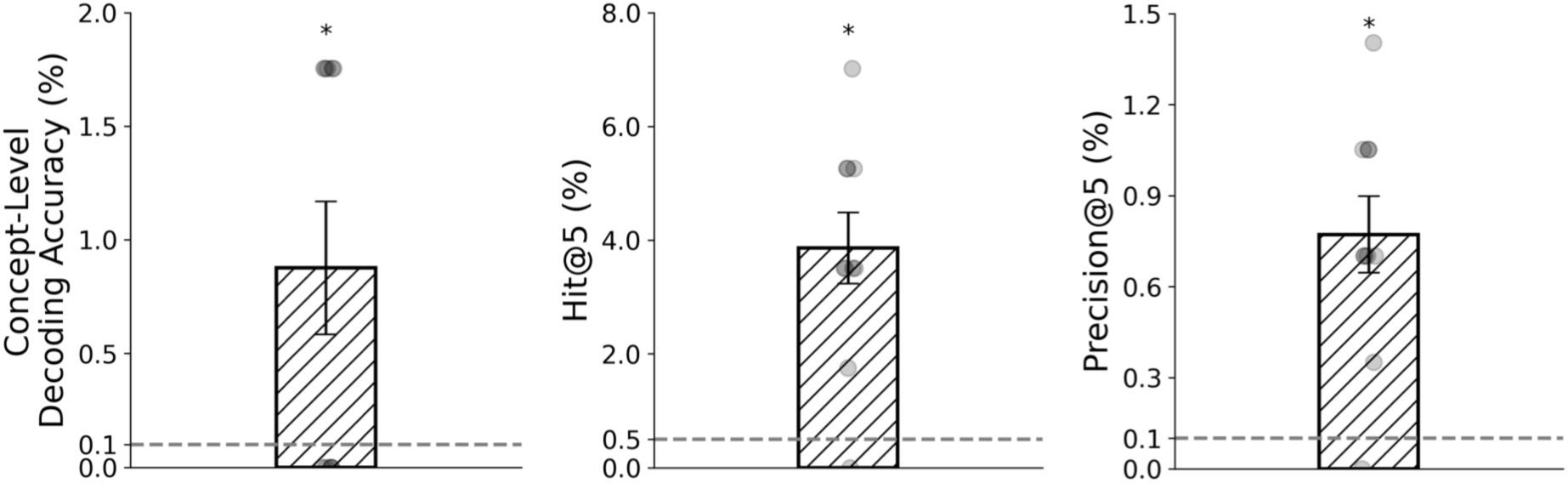
Database-based image reconstruction from EEG across the full ImageNet category space. This analysis extends the reconstruction shown in Figure 4B by expanding the candidate pool from the 57 ImageNet concepts shared with THINGS EEG2 to images spanning all 1,000 ImageNet categories. Real EEG responses corresponding to the 57 THINGS EEG2 test concepts that overlapped with ImageNet were used as queries against simulated EEG responses generated for ImageNet validation images spanning all 1,000 ImageNet categories. Candidate images were ranked according to the similarity between their simulated EEG responses and the measured EEG pattern. Reconstruction performance remained significantly above chance for Top-1 concept accuracy, Hit@5, and Precision@5 despite the substantially expanded candidate space. Bars indicate mean ± SEM across participants; asterisks denote significant above-chance performance (p < .05, FDR-corrected).

## Notes

### Competing Interest Statement

The authors have declared no competing interest.

## References

1. Bentin, S., Allison, T., Puce, A., Perez, E., & McCarthy, G. (1996). Electrophysiological Studies of Face Perception in Humans. Journal of Cognitive Neuroscience, 8(6), 551–565. 10.1162/JOCN.1996.8.6.551

2. Deng, J., Dong, W., Socher, R., Li, L. J., Li, K., & Fei-Fei, L. (2009). ImageNet: A Large-Scale Hierarchical Image Database. 2009 IEEE Conference on Computer Vision and Pattern Recognition, CVPR 2009, 248–255. 10.1109/CVPR.2009.5206848

3. Du, C., Fu, K., Li, J., & He, H. (2023). Decoding Visual Neural Representations by Multimodal Learning of Brain-Visual-Linguistic Features. IEEE Transactions on Pattern Analysis and Machine Intelligence, 45(9), 10760–10777. 10.1109/TPAMI.2023.3263181

4. Dube, B., Pidaparthi, L., & Golomb, J. D. (2022). Visual Distraction Disrupts Category-tuned Attentional Filters in Ventral Visual Cortex. Journal of Cognitive Neuroscience, 34(8), 1521– 1533. 10.1162/JOCN_A_01870

5. Finlayson, N. J., Zhang, X., & Golomb, J. D. (2017). Differential patterns of 2D location versus depth decoding along the visual hierarchy. NeuroImage, 147, 507–516. 10.1016/J.NEUROIMAGE.2016.12.039

6. Geirhos, R., Narayanappa, K., Mitzkus, B., Thieringer, T., Bethge, M., Wichmann, F. A., & Brendel, W. (2021). Partial success in closing the gap between human and machine vision. Advances in Neural Information Processing Systems 34, *34*, 23885–23899.

7. Gifford, A. T., Dwivedi, K., Roig, G., & Cichy, R. M. (2022). A large and rich EEG dataset for modeling human visual object recognition. NeuroImage, 264, 119754. 10.1016/J.NEUROIMAGE.2022.119754

8. Gifford, A. T., Lahner, B., Saba-Sadiya, S., Vilas, M. G., Lascelles, A., Oliva, A., Kay, K., Roig, G., & Cichy, R. M. (2023). The Algonauts Project 2023 Challenge: How the Human Brain Makes Sense of Natural Scenes. ArXiv. 10.48550/arXiv.2301.03198

9. Hebart, M. N., Dickter, A. H., Kidder, A., Kwok, W. Y., Corriveau, A., Van Wicklin, C., & Baker, C. I. (2019). THINGS: A database of 1,854 object concepts and more than 26,000 naturalistic object images. PLoS ONE, 14(10), 1–24. 10.1371/journal.pone.0223792

10. Hinojosa, J. A., Mercado, F., & Carretié, L. (2015). N170 sensitivity to facial expression: A meta-analysis. Neuroscience & Biobehavioral Reviews, 55, 498–509. 10.1016/J.NEUBIOREV.2015.06.002

11. Huth, A. G., De Heer, W. A., Griffiths, T. L., Theunissen, F. E., & Gallant, J. L. (2016). Natural speech reveals the semantic maps that tile human cerebral cortex. Nature, 532(7600), 453–458. 10.1038/nature17637

12. Huth, A. G., Nishimoto, S., Vu, A. T., & Gallant, J. L. (2012). A Continuous Semantic Space Describes the Representation of Thousands of Object and Action Categories across the Human Brain. Neuron, 76(6), 1210–1224. 10.1016/j.neuron.2012.10.014

13. Kingma, D. P., & Ba, J. L. (2014). Adam: A Method for Stochastic Optimization. 3rd International Conference on Learning Representations, ICLR 2015 - Conference Track Proceedings. https://arxiv.org/pdf/1412.6980

14. Kriegeskorte, N., & Douglas, P. K. (2019). Interpreting encoding and decoding models. Current Opinion in Neurobiology, 55, 167–179. 10.1016/J.CONB.2019.04.002

15. Kriegeskorte, N., Mur, M., & Bandettini, P. (2008). Representational similarity analysis - connecting the branches of systems neuroscience. Frontiers in Systems Neuroscience, 4. 10.3389/NEURO.06.004.2008

16. Kubilius, J., Schrimpf, M., Kar, K., Rajalingham, R., Hong, H., Majaj, N. J., Issa, E. B., Bashivan, P., Prescott-Roy, J., Schmidt, K., Nayebi, A., Bear, D., Yamins, D. L. K., & Dicarlo, J. J. (2019). Brain-Like Object Recognition with High-Performing Shallow Recurrent ANNs. Advances in Neural Information Processing Systems (NeurIPS), 32.

17. Kubilius, J., Schrimpf, M., Nayebi, A., Bear, D., Yamins, D. L. K., & DiCarlo, J. J. (2018). CORnet: Modeling the Neural Mechanisms of Core Object Recognition. BioRxiv. 10.1101/408385

18. Li, D., Wei, C., Li, S., Zou, J., & Liu, Q. (2024). Visual Decoding and Reconstruction via EEG Embeddings with Guided Diffusion. Advances in Neural Information Processing Systems, 37. 10.52202/079017-3266

19. Li, J., Li, D., Savarese, S., & Hoi, S. (2023). BLIP-2: Bootstrapping Language-Image Pre-training with Frozen Image Encoders and Large Language Models. Proceedings of the International Conference on Machine Learning (ICML), 19730–19742.

20. Lu, Z., & Wang, Y. (2025). Teaching CORnet Human fMRI Representations for Enhanced Model-Brain Alignment. Cognitive Neurodynamics, 19(1), 61. 10.1007/S11571-025-10252-Y

21. Lu, Z., Wang, Y., & Golomb, J. D. (2026). Achieving more human brain-like vision via human EEG representational alignment. Communications Biology, 9(1), 463-. 10.1038/s42003-026-09685-w

22. Luck, S. J., Heinze, H. J., Mangun, G. R., & Hillyard, S. A. (1990). Visual event-related potentials index focused attention within bilateral stimulus arrays. II. Functional dissociation of P1 and N1 components. Electroencephalography and Clinical Neurophysiology, 75(6), 528–542. 10.1016/0013-4694(90)90139-B

23. Mangun, G. R., & Hillyard, S. A. (1990). Allocation of visual attention to spatial locations: Tradeoff functions for event-related brain potentials and detection performance. Perception & Psychophysics, 47(6), 532–550. 10.3758/BF03203106

24. Mercure, E., Kadosh, K. C., & Johnson, M. H. (2011). The N170 shows differential repetition effects for faces, objects, and orthographic stimuli. Frontiers in Human Neuroscience, 5(JANUARY), 1–10. 10.3389/FNHUM.2011.00006

25. Nag, S., Berman, D., & Golomb, J. D. (2019). Category-selective areas in human visual cortex exhibit preferences for stimulus depth. NeuroImage, 196, 289–301. 10.1016/J.NEUROIMAGE.2019.04.025

26. Naselaris, T., Kay, K. N., Nishimoto, S., & Gallant, J. L. (2011). Encoding and decoding in fMRI. NeuroImage, 56(2), 400–410. 10.1016/J.NEUROIMAGE.2010.07.073

27. Naselaris, T., Olman, C. A., Stansbury, D. E., Ugurbil, K., & Gallant, J. L. (2015). A voxel-wise encoding model for early visual areas decodes mental images of remembered scenes. NeuroImage, 105, 215–228. 10.1016/J.NEUROIMAGE.2014.10.018

28. Pennington, J., Socher, R., & Manning, C. D. (2014). GloVe: Global Vectors for Word Representation. Empirical Methods in Natural Language Processing (EMNLP), 1532–1543.

29. Proverbio, A. M. (2021). Sexual dimorphism in hemispheric processing of faces in humans: A meta-analysis of 817 cases. Social Cognitive and Affective Neuroscience, 16(10), 1023–1035. 10.1093/SCAN/NSAB043

30. Radford, A., Kim, J. W., Hallacy, C., Ramesh, A., Goh, G., Agarwal, S., Sastry, G., Askell, A., Mishkin, P., Clark, J., Krueger, G., & Sutskever, I. (2021). Learning Transferable Visual Models From Natural Language Supervision. Proceedings of the International Conference on Machine Learning (ICML).

31. Rajalingham, R., Issa, E. B., Bashivan, P., Kar, K., Schmidt, K., & DiCarlo, J. J. (2018). Large-Scale, High-Resolution Comparison of the Core Visual Object Recognition Behavior of Humans, Monkeys, and State-of-the-Art Deep Artificial Neural Networks. Journal of Neuroscience, 38(33), 7255–7269. 10.1523/JNEUROSCI.0388-18.2018

32. Rossion, B., & Jacques, C. (2012). The N170: Understanding the Time Course of Face Perception in the Human Brain. In The Oxford handbook of event-related potential components (Oxford Uni, pp. 115–141). Oxford University Press. 10.1093/OXFORDHB/9780195374148.013.0064

33. Santos-Mayo, A., Gilbert, F., Mirifar, A., Tebbe, A.-L., Fang, R., Ding, M., & Keil, A. (2026). Concept2Brain: an AI model for predicting neurophysiological responses to text and pictures. Nature Communications. 10.1038/S41467-026-75653-X

34. Schrimpf, M., Kubilius, J., Hong, H., Majaj, N. J., Rajalingham, R., Issa, E. B., Kar, K., Bashivan, P., Prescott-Roy, J., Geiger, F., Schmidt, K., Yamins, D. L. K., & DiCarlo, J. J. (2020). Brain-Score: Which Artificial Neural Network for Object Recognition is most Brain-Like? BioRxiv. 10.1101/407007

35. Song, K., Tan, X., Qin, T., Lu, J., & Liu, T.-Y. (2020). MPNet: Masked and Permuted Pre-training for Language Understanding. Advances in Neural Information Processing Systems, 33, 16857–16867.

36. Song, Y., Liu, B., Li, X., Shi, N., Wang, Y., & Gao, X. (2024). Decoding Natural Images from EEG for Object Recognition. International Conference on Learning Representations (ICLR).

37. Stoinski, L. M., Perkuhn, J., & Hebart, M. N. (2023). THINGSplus: New norms and metadata for the THINGS database of 1854 object concepts and 26,107 natural object images. Behavior Research Methods, 1–21. 10.3758/s13428-023-02110-8

38. Wardle, S. G., Taubert, J., Teichmann, L., & Baker, C. I. (2020). Rapid and dynamic processing of face pareidolia in the human brain. Nature Communications, 11(1), 4518-. 10.1038/s41467-020-18325-8

39. Yamins, D. L. K., & DiCarlo, J. J. (2016). Using goal-driven deep learning models to understand sensory cortex. Nature Neuroscience, 19(3), 356–365. 10.1038/nn.4244

40. Zhang, K., He, L., Jiang, X., Lu, W., Wang, D., & Gao, X. (2025). CognitionCapturer: Decoding Visual Stimuli from Human EEG Signal with Multimodal Information. Proceedings of the AAAI Conference on Artificial Intelligence, 39(13), 14486–14493. 10.1609/AAAI.V39I13.33587

